# A Novel Optimization-Based Tool to Automate Infeasible Cycle-Free Gapfilling of Genome-Scale Metabolic Models

**DOI:** 10.1101/796706

**Authors:** Wheaton L. Schroeder, Rajib Saha

## Abstract

Stoichiometric metabolic modeling, particularly Genome-Scale Models (GSMs), is now an indispensable tool for systems biology. The model reconstruction process typically involves collecting information from public databases; however, incomplete systems knowledge leaves gaps in any reconstruction. Current tools for addressing gaps use databases of biochemical functionalities to address gaps on a per-metabolite basis and can provide multiple solutions, but cannot avoid Thermodynamically Infeasible Cycles (TICs), invariably requiring lengthy manual curation. To address these limitations, this work introduces an optimization-based multi-step method named OptFill which performs TIC-avoiding whole-model gapfilling. We applied OptFill to three fictional prokaryotic models of increasing sizes and to a published GSM of *Escherichia coli*, *i*JR904. This application resulted in holistic and infeasible cycle free gapfilling solutions. Part of OptFill can, in addition, be adapted to automate inherent TICs identification in any GSM, such as *i*JR904. Overall, OptFill can address critical issues in automated development of high-quality GSMs.

**In Brief:** Stoichiometric models of metabolism are useful in studying metabolic interactions in biological systems, but are labor-intensive to create, particularly when addressing gaps or cycles in metabolic reconstruction process. Introduced here is a novel tool, OptFill, which can be used to address both gaps and cycles in model reconstruction, increasing automation.

**Highlights:**

- This work presents an alternative to state-of-the-art methods for gapfilling.
- Unlike current methods, this method is holistic and infeasible cycle free.
- This method is applied to three test and one published model.
- This method might also be used to address infeasible cycling.

## INTRODUCTION

The use of systems biology in uni- and multi-cellular organisms to engineer or enhance desirable phenotypes and study system-wide metabolic processes in microbes, plants, and animal systems, is well-established and capable of affecting the lives of millions of individuals, such as in the case of artemisinin production in yeast or enhancing the nutritional value of agricultural products (Beyer *et al*., 2002) (Hall, Brouwer and Fitzgerald, 2008). As opposed to traditional qualitative approaches, computational approaches based on stoichiometric Genome-Scale Models (GSMs) of metabolism can be used to predict non-intuitive genetic interventions (Srinivasan, Cluett and Mahadevan, 2015) by accounting for gene-protein-reaction (GPR) links. GSMs may also lead to increased understanding of how a change in environment, a change in organism nutrition, or a gene knockout, can affect the entire metabolic system of an organism or community through tools such as Flux Balance Analysis (FBA) (Orth, Thiele and Palsson, 2010), OptKnock (Burgard, Pharkya and Maranas, 2003), and OptForce (Ranganathan, Suthers and Maranas, 2010). GSMs have been developed for many prokaryotic (Magnúsdóttir *et al*., 2016)(Shoaie *et al*., 2013), animal (Brunk *et al*., 2018), plant (Gomes de Oliveira Dal’Molin *et al*., 2015)(Saha, Suthers and Maranas, 2011), and fungal (Andersen, Nielsen and Nielsen, 2008)(Liu *et al*., 2013) systems, enhancing mechanistic understanding and exploration of system-wide metabolism in such organisms as *E. coli* (Ranganathan, Suthers and Maranas, 2010), cyanobacteria (Saha *et al*., 2016), yeast (Ng *et al*., 2012), and other species (Saha, Suthers and Maranas, 2011)(Gudmundsson, Agudo and Nogales, 2017)(Shoaie *et al*., 2013). GSMs are typically reconstructed by gleaning information on gene annotations, enzyme functions, associated reactions, and reaction directionality from major public databases such as KEGG (Kanehisa *et al*., 2017), ModelSEED (Overbeek *et al*., 2005), the NCBI (Limviphuvadh *et al*., 2018), MetaCyc (Caspi, 2006), K-Base (Arkin *et al*., 2018), and BIGG (King *et al*., 2016). At present, there is no complete knowledge of any genome. For instance, the annotated genome of one of the most prolifically studied organisms, *Escherichia coli* strain K-12 substrain MG1655, contains about 6.8% putative proteins and 16.1% uncharacterized proteins (UniProtKB, 2018). Furthermore, approximately 61% of proteins lack an Enzyme Commission (EC) number, which is important for the identification of GPR links in any GSM reconstruction (UniProtKB, 2018). Inevitably, incomplete gene annotation and system knowledge (including reaction direction) leaves metabolic gaps, imbalances, or Thermodynamically Infeasible Cycles (TICs) in any initial GSM reconstructions, leaving the model incomplete. Particularly problematic are TICs, sets of reactions which can carry flux in the absence of nutrition provided to the model because their net stoichiometry is zero, also known as futile cycles or type III reactions (Thiele and Palsson, 2010). These cycles can negate metabolic costs (Thiele and Palsson, 2010), report infeasibly large reaction rates, are difficult to identify (De Martino *et al*., 2013)(Schellenberger, Lewis and Palsson, 2011), and can inhibit the proper function of optimization-based tools which rely on duality to optimize multiple objectives such as OptKnock (Burgard, Pharkya and Maranas, 2003) and OptForce (Ranganathan, Suthers and Maranas, 2010).

A significant limitation on the time required to reconstruct GSMs is the amount of time and manual labor required to curate these incomplete reconstructed model, addressing various issues such as element and charge balances; reaction directionality; metabolic gaps; TICs; and other inconsistencies. Addressing these issues is time and labor intensive, often requiring months to years of manpower before a predictive model is generated (Thiele and Palsson, 2010), which is a prerequisite for conducting research on phenotypic enhancement or study metabolism. Two of the most challenging aspects of model development are the identification and elimination of TICs and the resolving of metabolic gaps.

The existing methods/tools that have been developed to address the identification and resolution of TICs can be broadly categorized into two groups: first, methods that can identify existing TICs in a model (De Martino *et al*., 2013), and second, methods that can force no-flux through existing TICs in a model (Schellenberger, Lewis and Palsson, 2011)(Nigam and Liang, 2007). Although developing these is a significant step toward building a better and more predictive GSM, there remain challenges that need to be addressed. For example, the Monte Carlo sampling-based method (De Martino *et al*., 2013) cannot guarantee the identification of all TICs. The second approach is the avoidance of TICs by the application of Kirchoff’s Loop Law in methods such as Loopless COBRA (Schellenberger, Lewis and Palsson, 2011). This approach does successfully avoid TICs, but does not address the root cause in the model which can make some models problematic for tools such as OptForce which require no TICs (Ranganathan, Suthers and Maranas, 2010). Another approach is the addition of thermodynamic constraints to the model using known thermodynamic quantities (Nigam and Liang, 2007), which works well for well-studied organisms for which these *in vivo* parameters are known, but is more difficult to implement for non-model organisms.

To address and resolve metabolic gaps, GapFind and GapFill (Satish Kumar, Dasika and Maranas, 2007) are some of the most common tools used (Pitkänen *et al*., 2014)(Henry *et al*., 2010)(Kim *et al*., 2012). GapFind and GapFill are optimization-based Mixed Integer Linear Programming (MILP) problems, and have been successfully implemented in the reconstruction of metabolic models, of prokaryotic and eukaryotic biological systems such as cyanobacteria (*Synechocystis* sp. PCC 6803 and *Cyanothece sp ATCC 51142*) (Saha *et al*., 2012), corn (*Zea mays*) (Simons *et al*., 2014), yeast (*Saccharomyces cerevisiae*), and Chinese hamster ovary cells (Chowdhury, Chowdhury and Maranas, 2015). Other methods of automated gapfilling which build on the capabilities of GapFill include GenDev (Latendresse and Karp, 2018), FastDev (Latendresse and Karp, 2018), likelihood-based gapfilling (Karp, Weaver and Latendresse, 2018), and phenotype-based gapfilling (Cuevas *et al*., 2019). All these tools are constructed with the aim of increasing the accuracy of the GapFilling method, through comparison to some level of data such as phylogenetic, phenotypic, or genetic. In this work, a problematic aspect of all these tools is considered which these other tools were not built to address. Despite their success, the tools for gapfilling have significant limitations including: i) gaps are addressed on a per-metabolite basis (as opposed to a whole-model holistic approach), ii) thermodynamic feasibility is often not considered, and iii) reaction direction is not considered in gapfilling, rather all reactions are added reversibly. From the first and second limitations, several problems arise including: i) inability to guarantee that the minimum number of reactions are added to fix metabolic gaps on a whole-model basis; ii) inability to identify and avoid unfavorable interactions between multiple gap fixes (often, TICs); and iii) differences resultant model dependent on the individual curator.

To address current TIC-finding and gapfilling limitations, this work introduces a multi-step optimization-based MILP method. The first step is to solve an iterative optimization-based TIC-Finding Problem (TFP) which identifies potential TICs, which may be caused by adding reactions from a database in a given direction (see Figure 1). This problem is unique in that it considers the direction of reactions participating in TICs and can be easily adapted for the purposes of model curation sans database for the resolution of inherent TICs. The second step involves the solving of three optimization-based problem, the Connecting Problems (CPs), which are highly similar but have different objectives. The first Connecting Problem (CP1) is the maximization of model metabolites successfully connected to metabolic network, e.g. maximizing the number of metabolites which the connected model can now produce, while avoiding the addition of TICs. The second Connecting Problem (CP2) is the minimization of the number of reactions required to achieve the objective of CP1. The third connecting problem (CP3) is the maximization of the number of reactions to be added reversibly from the database to achieve the objectives of CP1 and CP2 subject to avoiding TICs. As proof of concept, the OptFill approach is applied to three test stoichiometric models of increasing sizes (models of 28 to 210 reactions, databases of 17 to 77 reactions) with designed metabolic gaps, and one smaller (1074 reactions) GSM of *Escherichia coli* with acknowledged metabolic gaps (Reed *et al*., 2003) using another GSM of *E. coli* as the basis for a database (Feist *et al*., 2007a). With the computational resources at hand, the full OptFill method is limited to relatively smaller stoichiometric models and databases but should be applicable to larger models and databases given access to greater computational power.

**Figure 1:**
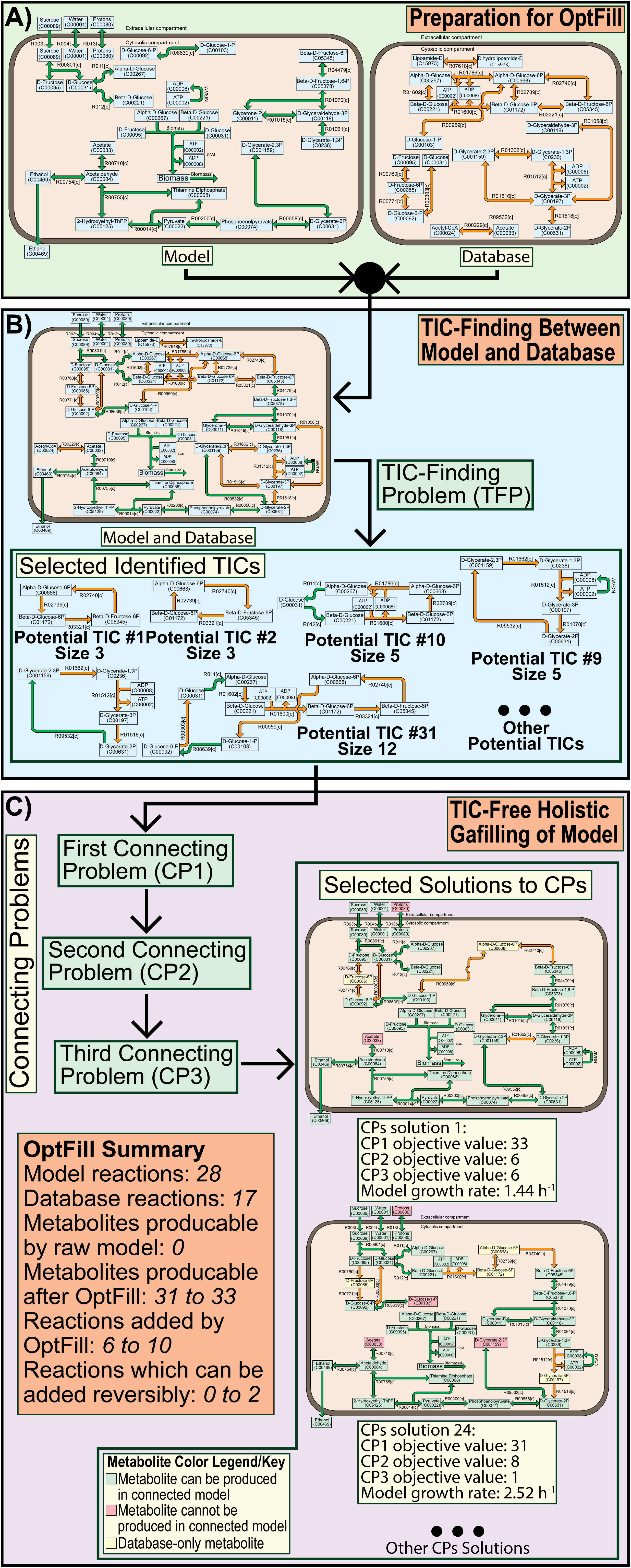
Visualization of OptFill results with respect to the first test model and database. Background colors of this visualization correspond to the workflow presented in Figure 2, where the colors green, blue, and purple correspond to preparation for OptFill; the TFP and its framework; and the CPs and its framework, respectively in both images. A) Shows that the model and database are separate, but are both used in the workflow to prepare for OptFill and in OptFill itself. B) Shows how the combined model and database might appear, and how this combination is used in the TIC-Finding Problem to identify potential TICs which might occur between the model and the database. Selected identified potential TICs are shown here as illustrative examples. Potential TICs #1 and #2 illustrate how TICs occurring in different directions are identified as separate TICs, how identified TICs might only occur between database reactions, and illustrate the two of the smallest identified TICs. Potential TICs #9 illustrates a larger TIC which makes use of an irreversible reaction (NGAM), and therefore has no opposite-direction TIC, making the direction of the other reactions important. Potential TICs #10 and #31 illustrate infeasible cycling involving an energy molecule (ADP/ATP), in addition to potential TIC #31 being the largest identified TIC. C) Show the application of the Connecting Problems (CPs) and the first and last solution of the CPs. These solutions differ in the number of model metabolites which could not be connected (red boxes); the number of new metabolites introduced to the model (yellow boxes); the number and reversibility of database reactions added (orange arrows); and the resultant model growth rate.

## RESULTS

### Development of OptFill

OptFill was conceived and developed to address the limitations of the current state-of-the-art GapFind/GapFill (Satish Kumar, Dasika and Maranas, 2007) tool. The initial stages of the design-build-test (DBT) cycle, involving the first Test Model (TM1) and the first Test Database (TDb1) involved only a single connecting problem. TM1 was constructed as a small stoichiometric model involving starch metabolism and glycolysis to produce ethanol but with metabolic gaps preventing growth (see Figure 1). TDb1 was designed to have the capacity to fill these gaps, at the expense of potentially producing TICs. In the DBT cycle, it was soon realized that the TFP was necessary to define the potential TICs which might occur. The TFP was built to solve for the smallest TICs first, that is, the TIC with the smallest number of participant reactions, and then solve for larger TICs to prevent multiple TICs masquerading as a single TIC solution. The workflow surrounding the TFP is shown in Figure 2. The CPs were developed to ensure consistency in the number, order, and identity of the CP solutions while avoiding the addition of whole TICs identified as potentially occurring between the model and database. See Figure 3 for the conceptual formulation of each type of problem. All problems which are part of the OptFill tool are Mixed Integer Linear Programming (MILP) problems which ensure global optimality of each solution in each iteration.

**Figure 2:**
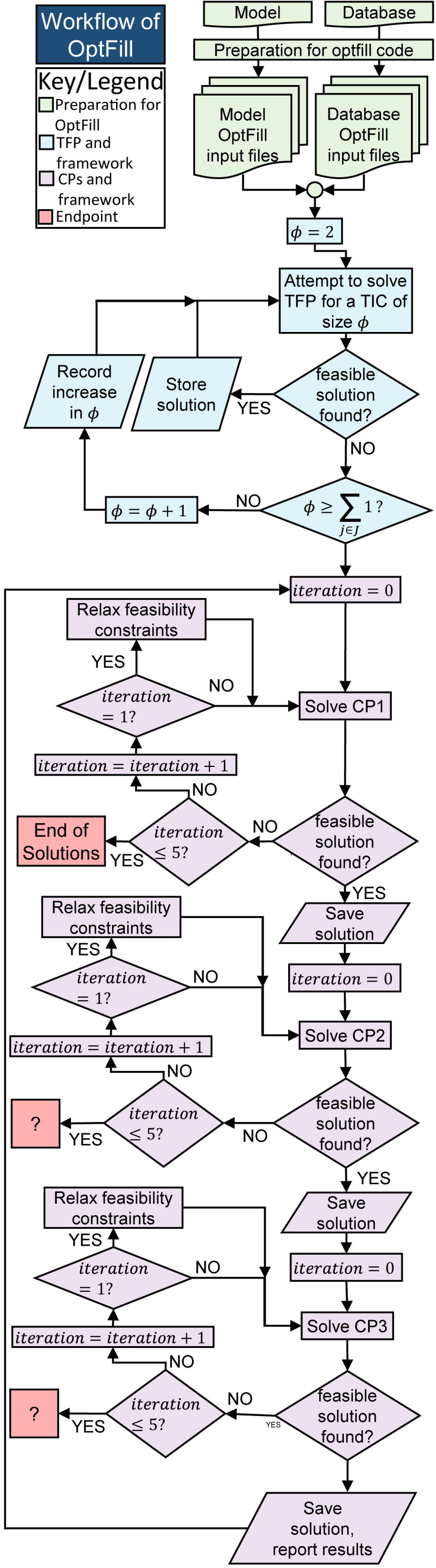
Workflow of OptFill. This is a workflow diagram of the OptFill tool. Green nodes represent the preparatory workflow, blue the workflow of the TIC-Finding Problem (TFP), brown the workflow of the Connecting Problems (CPs), purple the error-handling workflow imbedded in the CPs workflow, and red the endpoints of the workflow. This color scheme is consistent with Figure 1. It should be noted that only one endpoint truly exists, when a solution to CP1 is not found, because the other problems, CP2 and CP3 will have solutions if CP1 has a solution, hence the workflow exit points being represented by a question mark at these points.

**Figure 3:**
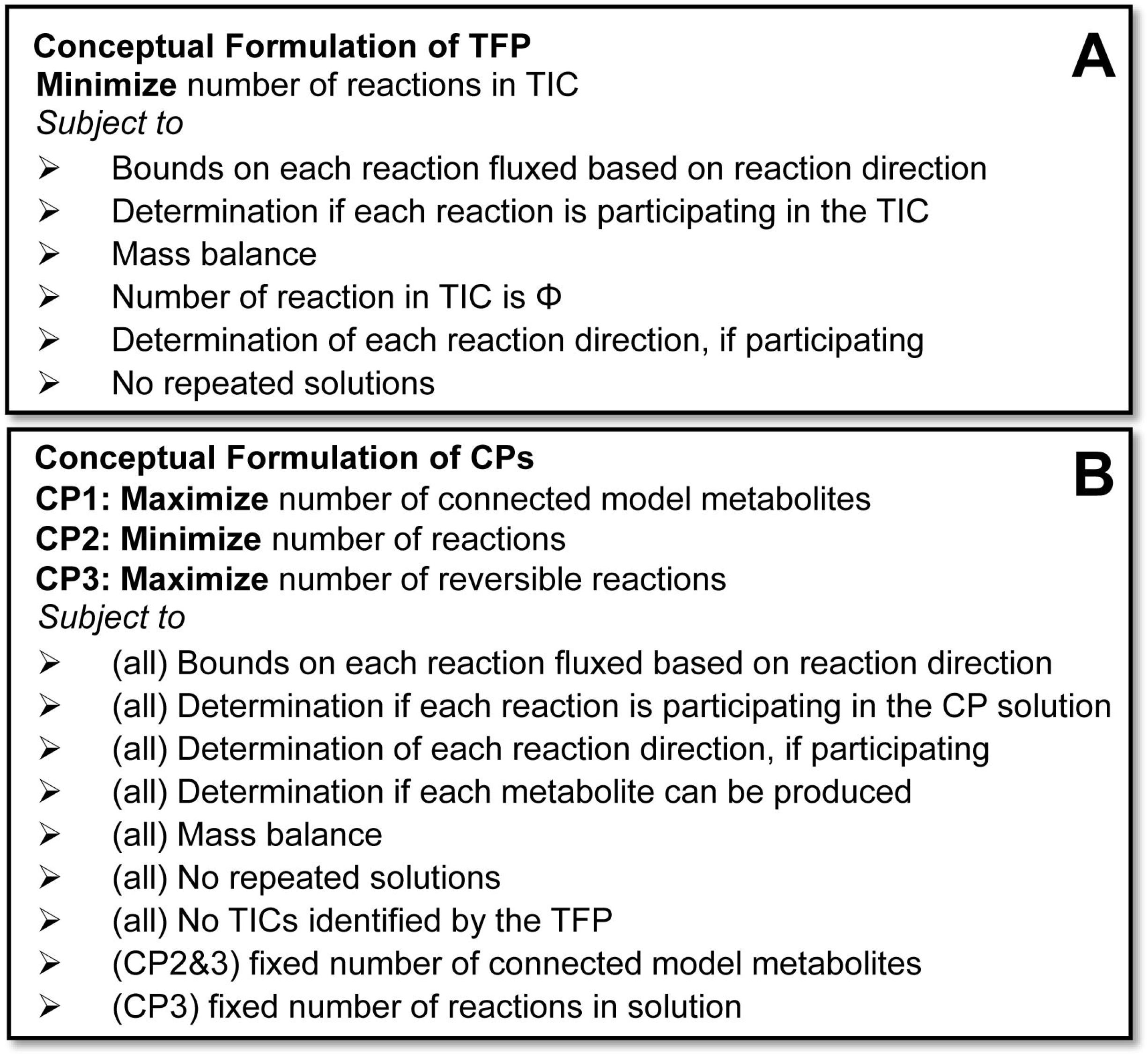
Conceptual formulation of each problem in OptFill. This figure give a conceptual formulation of the TIC-Finding Problem, TFP, in part (A) and the Connecting Problems, CP1, CP2, and CP3, in part (B). In part (B), as three connecting problems are solved, each conceptual constraint has indicated CPs to which it is applied. Conceptual constraints may require multiple mathematical constraints to be realized, see methods for mathematical formulation.

On occasion, the feasibility constraints used may be too strict to return a feasible solution to the CP problems, this occasionally resulted in execution errors prematurely ending OptFill before completion. Therefore, an error handling framework was built around each CP problem allowing a one-time relaxation of feasibility constraints for each problem. These frameworks are shown in Figure 2. OptFill is ended when CP1 no longer has a feasible solution even when feasibility constraints are relaxed (which occurs because previous solutions are prevented from being re-identified) since at that point none of the CP2 and CP3 will have a feasible solution. Further, all OptFill runs described used nonstandard CPLEX solver options, which effectively eliminated most types of cuts which cause some level of reduction to the solution space, particularly those which could result in non-optimal solutions being reported as optimal. These include flow, zero-half, and Gomory fractional cuts among others. This was done because the order of solutions is important in the OptFill method, and the order of solutions also has bearing on the number of solutions returned. See methods for further detail.

### Application of OptFill to Test Models

After finalizing the formulation (Figure 3 and methods) and workflow (Figure 2) of OptFill, a detailed analysis of OptFill results with respect to TM1 and TDb1 was undertaken. Some qualitative results of the application of the OptFill workflow to TM1/TDb1 are shown in Figure 1, which include the initial model and database, Figure 1(A); the combination of the model and database, Figure 1(B); selected identified potential TICs, Figure 1(B); and selected identified CPs’ solutions, Figure 1(C). As is shown in Figure 1(A), TM1 is too disconnected to produce biomass, but in combination with TDb1 can potentially produce biomass. When the TFP is applied (Figure 1(B)) 31 potential TICs consisting of 3 to 12 reactions (hereafter, sizes 3 to 12) were identified. The average solution time (when a solution was found) is 0.175 s (σ=0.0727 s, min=0.0870 s, max=0.378 s). It should be noted that all solve times reported here are not constant, even if using same resources. Figure 1 (B) highlights 5 potential TICs which were identified. The first two TICs identified, TIC #1 and #2, show that the TFP can identify TICs occurring only in the database; that TICs consisting of the same metabolites and reactions are identified separately if reaction directions are different; and shows two of the smallest TICs identified. Potential TIC #9 shows a TIC of moderate size (for TM1/TDb1) which contains an irreversible model reaction related to Non-Growth Associated Maintenance (NGAM) and therefore will not have a companion potential TIC of opposite direction, unlike potential TIC #1 and TIC #2 (in the opposite direction). Further, this highlights the potential for infeasible cycling which effectively negates the cost of NGAM of the model. If added in its entirety, NGAM would be irrelevant to the model at any value and would significantly reduce model accuracy. This TIC might not be manually identified since NGAM is usually a fixed quantity. Potential TIC #10 highlights another type of infeasible cycling involving ADP/ATP, but this cycling essentially negates the cost of phosphorylation/ dephosphorylation of glucose-6-phosphate isomers. Finally, potential TIC #31 is included to highlight a non-intuitive identified TIC, in addition to being the largest TIC identified. This TIC involves the separate cycling of sugars and 3-carbon molecules linked and is made possible by ADP/ATP cycling (sugar cycling consumes ATP and 3-carbon cycling produces ATP). These examples illustrate that many, but not all, potential TICs involve the infeasible cycling of energy molecules, which should be particularly avoided in the reconstruction of models of metabolism as this can result in negated costs for various biological activities with which a cost should be associated. This negated cost can often result in increased model growth rate and reaction fluxes, reducing the model’s accuracy.

The model, database, and TFP solutions form the input for the CPs. Before solving the CPs, a modified version of CP1 was run which prohibited the addition of database reactions. This modified CP1 reported that the raw TM3 model was capable of producing no metabolites. The CPs, when applied to TM1 and TDb1, identified 24 potential solutions which connected between 31 and 33 metabolites with the additions of 6 to 10 reactions, of which 0 to 6 could be reversible without TICs. The average time to solve all three CPs for each solution was 0.639 s (σ=0.147 s, min=0.433 s, max=0.950 s), see Figure 4. From the FBA performed on each connecting problem solution with the objective of maximization of biomass, the mean maximum biomass production rate of the set of connected models was 2.43 h^−1^ (σ=0.394 h^−1^, min=1.44 h^−1^, max=2.90 h^−1^). Solution times for the FBA code were not recorded as FBA solution time is generally low. Two connecting problem solutions, the first and the last, are shown in Figure 1(C). These solutions are notably different in the number of model metabolites connected by the CPs’ solution (green boxes in the metabolic sketch), the number of intermediate metabolites introduced by these solutions (yellow boxes), the number of database reactions introduced (orange arrows), and even the use of energy molecules. For instance, CPs’ solution 1 introduces only two additional metabolites and 6 reactions reversibly from the database which were part of the CPs’ solution and connects all but two model metabolites. The first is acetate, which is a dead-end metabolite. The second is the extracellular proton, which suggests that the model is small enough that all protons produced are also consumed. This solution had the slowest growth rate of all connecting problem solutions. On the other hand, CPs’ solution 24 connects two fewer metabolites than CPs’ solution 1, requires two more reactions, introduces two more intermediate metabolites, and has a higher growth rate. It is hypothesized that this is due to the more efficient production of ATP allowed by reaction R01512[c] (enzyme ATP:3-phospho-D-glycerate 1-phosphotransferace in the cytosol), which is present in many other high-biomass solutions. This reaction allows two dephosphorylation events to produce ATP, as opposed to only one (the other event occurring by hydrolysis).

**Figure 4:**
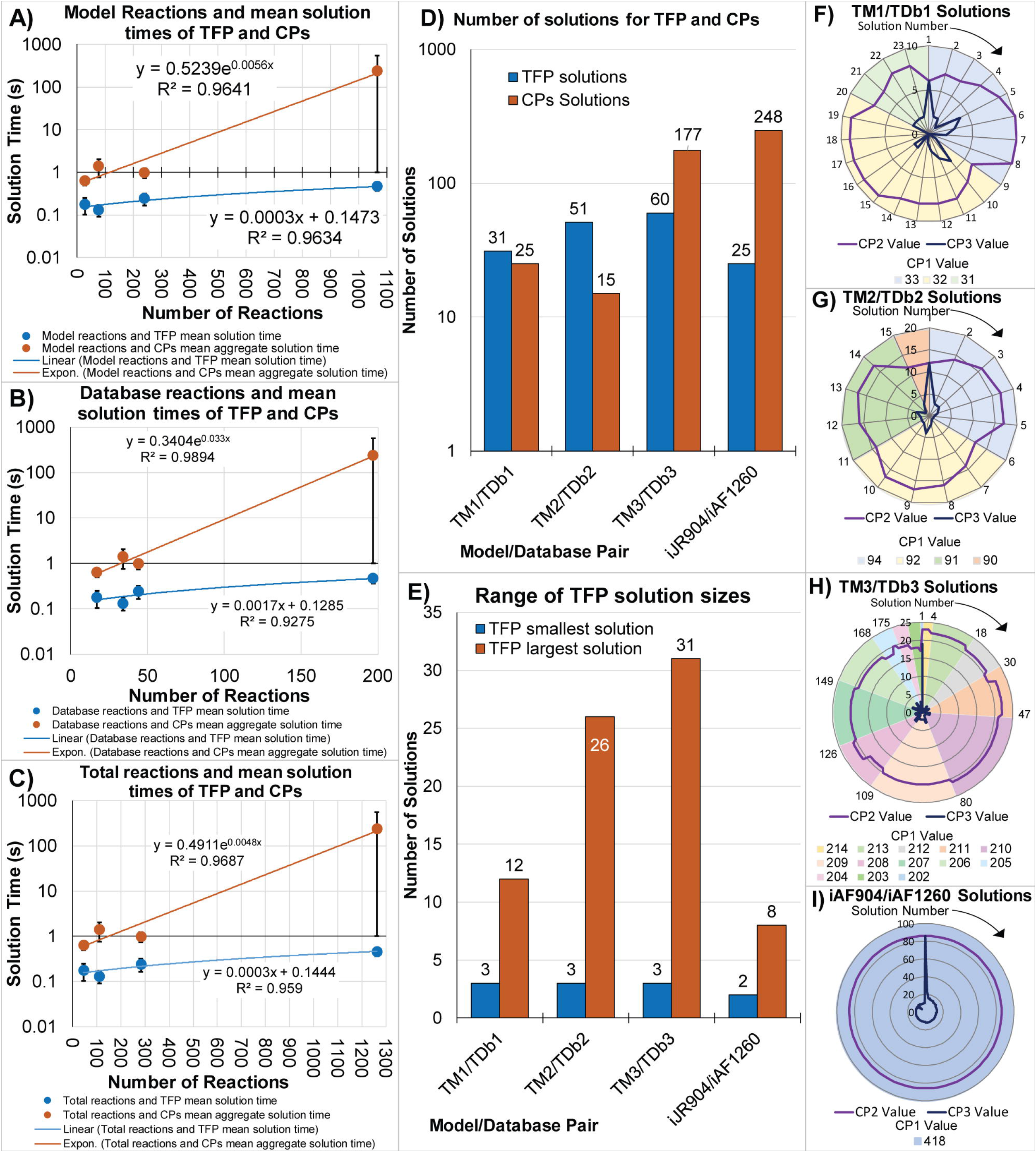
Visualization of OptFill solution time and results. This figure show the trends in solution time, (A) through (C), of the TIC-Finding Problem (TFP, blue) and the Connecting Problems (CPs, brown) with trend lines with the highest Pearson’s correlation coefficient of linear, exponential, power, and logarithmic fits. These trends are considered with respect to the number of reactions in the model (A), database (B), and total reactions (C). Parts (D) through (F) highlights the trends of solutions. Part (D) highlights the number of solutions found by the TFP and CPs; Part (E) highlights the range in size of the identified potential TICs by the TFP. Parts (F) through (I) highlight the variety of CPs’ solutions. In these figures, the pie-chart indicates the number of metabolites connected by the CP1 solution, and the radar chart is used to indicate the CP2 solution (number of reactions added) and the CP3 solution (number of those reactions that are added reversibly).

It was decided to build two larger test models to study the increase in number of solutions and time required to reach those solutions by OptFill to investigate its scale-up potential. Each test model was built from an OptFill solution of a previous solution to highlight the ability of this tool to be applied in sequence. In the application of OptFill to the existing models of organisms, careful attention must be paid in selection of a CPs’ solution to accept, including considerations of energy metabolism, predicted growth rates, and remaining unconnected metabolites. Here, CPs’ solution 1 was selected and combined with TM1 as the base of the second test model (TM2). Reactions and metabolites from the fatty acid biosynthesis and the pentose phosphate pathway were added to this base, in which gaps were manually created. Reactions which could address these gaps formed the second test database (TDb2). Redundant metabolic functions were added to TDb2 to allow for potential TICs. Similarly, the third test model (TM3) was built from the first CPs’ solution of TM2 and TDb2. Additionally, a bank of reactions from the amino acid synthesis pathways, including redundant functionalities, was created. This bank was automatically (randomly) sorted between those reactions which would be added to complete TM3 (~80% of new reactions) and those which would constitute the third test database (TDb3, ~20% of new reactions). As random sorting was used, a modified version of the TIC-Finding Problem (modified TIC-Finding Problem, mTFP), was used to identify inherent TICs in TM3 and TDb3 which resulted from the random assortment of new reactions. The reactions most-commonly participating in identified inherent TM3 TICs were moved to the TDb3 until no inherent TICs remained (5 reactions in total).

For OptFilling of TM2/TDb2, 51 TICs consisting of 3 to 26 reactions were identified by the TFP, with a mean solution time of 0.131 s (σ=0.0405 s, min=0.0850 s, max=0.308 s). The largest TIC, potential TIC #51 consisting of 26 reactions, would have largely been very difficult to identify by a non-automated method as it spans six KEGG pathways including glycolysis; the pentose phosphate pathway; purine metabolism; nicotinate and nicotinamide metabolism; starch and sucrose metabolism; riboflavin metabolism. TIC #51 involves the cycling of 3-, 4-, 5-, and 6-carbon molecules, energy molecules (ATP, NADH, and NADPH), and energy molecule hydrolysis.

Before solving the CPs, the modified CP1 was run and reported that the raw TM2 model was capable of producing no metabolites. Fifteen potential CPs’ solutions were identified which each connected 90 to 94 metabolites using 17 to 23 reactions, of which 0 to 19 could be reversible without TICs. The average time to solve all three CPs for each solution was 1.40 s (σ=0.639 s, min=0.404 s, max=2.65 s), see Figure 4. From the FBA performed, the biomass production rate of most CPs’ solutions applied to TM2 was 1.31 h^−1^, for 10 solutions, and 1.36 h^−1^ for the remaining five. In the CPs’ solutions, those with the highest biomass had fewer metabolites which could be connected (all solutions with higher biomass production were generated after lower biomass production solutions). Those with the higher biomass production rates generally had one fewer reaction which required ATP hydrolysis, and therefore had slightly more energy in the system to spend on the production of biomass that their lower biomass counterparts.

Similarly, OptFill applied to TM3/TDb3 resulted in the identification of 60 TICs consisting of 3 to 31 reactions by the TFP and 177 potential CPs’ solutions which each connected 202 to 214 metabolites with 12 to 17 reactions, of which 1 to 12 could be reversible without TICs. As before, the modified CP1 was used to identify 54 metabolites which the raw TM3 was capable of producing. The mean TFP solution time was 0.240 s (σ=0.0756 s, min=0.141 s, max=0.541 s), whereas the mean CPs’ solution time was 0.985 s (σ=0.249 s, min= 0.573 s, max=1.86 s), From the FBA performed, the mean biomass production rate of the connected model was 3.29 h^−1^ (σ=0.179 h^−1^, min=3.11 h^−1^, max=3.47 h^−1^). Runtime and solution metrics for all solutions are shown in Figure 4. Unlike TM1 and TM2 OptFilling results, there was no solution where all database reactions to be added by the CPs’ solution could be added reversibly. This indicates that, for all solutions, the direction in which database reactions are added is important to avoid TICs to produce a model without the disadvantages of TICs described previously. Furthermore, the biomass production rate did not appear as dependent on either the number of metabolites connected or reactions added as in previous CPs’ solution sets. Instead, the biomass production rate seems to most depend on the method of sulfate assimilation.

### Application of OptFill to *i*JR904

In order to show how the OptFill workflow might scale up to a GSM, the *i*JR904 model, consisting of 761 metabolites, 1074 reactions, and 904 genes, which models *Eschericia coli* (Reed *et al*., 2003) was selected as the base model to fix. The *i*AF1260 model, a model extending onto *i*JR904, consisting of 1598 metabolites, 2,381 reactions, and 1260 genes (Feist *et al*., 2007b) was selected to serve as the set of reactions from which to build the database. *i*JR904 contains 70 dead-end metabolites (Reed *et al*., 2003) which need fixing. Before applying OptFill, some minor formatting changes were made (described in methods and Supplemental File 1) and it was decided that carbon-limited aerobic growth using acetate would be the condition for which *i*JR904 model would be fixed. Metabolite exchange rates were taken from Reed *et al*., 2003 to describe this growth condition.

In order to create the database which would be applied to *i*JR904, all *i*AF1260 exchange reactions and reactions with names identical to those in *i*JR904 (which are assumed to be the same reaction as the former was built from the latter) were removed from *i*AF1260 to form the initial database which consisted of 1441 reactions. This proved too computationally intensive for the resources, and therefore this database was further simplified in a manner which it is suggested others with limited computational resources might also use. First, the *i*AF1260-based database and *i*JR904 were combined in single model file and Flux Variability Analysis (FVA) (Gudmundsson and Thiele, 2010) was performed (see Supplemental File 2). Those *i*AF1260 reactions capable of holding flux as determined by FVA (715 reactions) were defined as the new database.

OptFill was performed on *i*JR904 using this new database. This still resulted in a slow OptFill process, therefore solutions which were reported (4 identified in the allotted solve time of 24 hours) were collected. All *i*AF1260 reactions which participated in at least one solution (a total of 182 reactions) were selected as the basis of the third *i*AF1260-based database. This resulted in significantly lower computational requirements for the application of OptFill. This database proved, upon application of OptFill, to be without TICs. For the purposes of demonstration and showing how the increase of TFP solution time changes with model and database size, it was arbitrarily decided to add six reactions manually added from the previous database which could participate in potential TICs between the model and database, but which did not create TICs only within the database. Further, the mTFP was applied to the *i*JR904 model. From the mTFP results, it was noticed that in *i*JR904, some reactions were included in the model twice, both as reversible and irreversible, causing inherent TICs in the *i*JR904 model involving these duplicate reactions. It was decided to move the irreversible reactions of each duplicate pair to the database (nine reactions in total) so that all *i*JR904 models were still present in the OptFill in some capacity. The final *i*AF1260-based database for the OptFilling of *i*JR904 totals 188 reactions. Initial, final, and intermediate *i*AF1260-based databases used can be found in Supplemental Files 2 and 3.

Demonstrated here is a procedure by which the database applied to a model can be significantly decreased in size to reduce computational cost of the OptFill method, while still effectively addressing metabolic gaps. This can be summarized as i) eliminate all duplicate reactions; ii) perform FVA on a pseudomodel which is a combination of the database and model and use the results to eliminate reactions which cannot carry flux; and iii) perform OptFill using databases with larger solution time, collect a few sample solutions, and use the set of reactions participating in sampled solutions as the database. Applications of steps i) and ii) as well as iterative applications of iii) might be used by modelers to shrink the databased used in OptFilling to a size which is possible to solve in a modest period of time given the computational resources available.

This final *i*AF1260-based database was used to OptFill *i*JR904 model. In this final iteration, there were 25 TICs of size 2 to 8 reactions identified. The associated mean TFP solution time was 0.410 s (σ=0.0978 s, min= 0.330 s, max=0.687 s). The TICs identified were generally simple, as they stem from reactions manually added to the database which cause TICs. 11 TICs occur between just two reactions, and a further 4 involving only a single database reaction. Each of these effectively precluded a single database reaction from being added in a certain direction. When the CPs were applied to *i*JR904, it was found that the CPs’ solution time had increased considerably from that of other models, to a mean of 236 s (σ=329 s, min= 15.3 s, max=1010 s). The solution time of this model was significantly increased due to disabling of many types of cuts which a solver might use to decrease solution time, but which lead to non-optimal solutions being reported as optimal. These are particularly relevant because minor cuts, such as those that accept a 0.5% reduction in the optimal solution value, can change the number of metabolites connected by the CPs by two or more for GSMs. As the order of solutions is important, even these minor relaxations were deemed problematic and were therefore mostly disabled, leading to increased solution time. If these cuts were allowed, CPs’ solution time would have been approximately an order of magnitude less than reported here. The modified CP1 problem reported that the *i*JR904 model was capable of producing 358 metabolites under the given aerobic growth on acetate conditions, and all CPs’ solutions connected 418 metabolites with the addition of 86 reactions. All CPs’ solutions produced biomass at a rate of 0.108 h^−1^. This is likely a result of the database reduction steps taken. The variation on the CPs’ solution occurred in the number of connecting reactions which could be added reversibly, ranging from 5 to 86. It can be seen in Figure 4(I) that efforts to prevent non-optimal solutions from being reported as optimal were not entirely successful. There exists one CPs’ solution, solution #72, where the optimal (maximum) CP3 solution value is 5, whereas the optimal (maximum) CP3 solution value was 11 from solutions #71 and #73. This occurred when all solutions were subject to approximately the same constraints (save the integer cuts necessary to prevent repeated solutions). It is noted earlier that many types of cuts were disabled, but not all, and one type of cut or other solver setting allowed this non-optimal solution to be reported as optimal, however; eliminating all such cuts and settings proved prohibitively time-consuming. Therefore, the settings that can be found in Supplemental File 3 were selected as those which, for this work, best balanced solution order and solution time.

In the OptFilling solutions of *i*JR904, several trends can be noticed which were not present in the smaller test models. First, when performing FBA, with the objective of maximizing biomass, on the resultant OptFilled *i*JR904 model, not all reactions from the database held flux. This is because these reactions made it possible for the model to produce metabolites which are not required for the production of biomass or provides an alternative pathway for the production of biomass which might be less efficient. This does not mean that these connected metabolites are unimportant under other, equally valid, objective functions, for instance the connected metabolites may be bioproduction targets. Further, some TICs exist between *i*JR904 model reactions in the OptFill solutions and notably one database reaction. For most model reactions, these TICs occur because forward and reverse reactions are written separately. The TIC involving the database reaction resulted from the proton uptake exchange reaction being allowed a very high reaction flux in the *i*JR904 model. The TFP was performed with all exchange reactions fixed to a flux of zero, therefore the TFP did not identify this TIC which involved an exchange reaction. When the exchange reactions were allowed to carry flux again in the CPs, the high proton uptake rate (here, 1000 mmol/gDW·h) allowed the cycling of reactions. These resulting TICs highlighted two important considerations in using OptFill. First, the mTFP should be used in combination with manual editing of the model to ensure that the model does not contain inherent TICs as the usual OptFill workflow will not address inherent TICs. Second, reasonable bounds should be applied to all exchange reactions (such as the proton uptake reaction) and to forward and reverse reaction pairs to prevent TICs in the OptFilled model.

### OptFill Solution Times

With the caveats of the available resources (see the methods section for information on the software and hardware tools available for this work), the TFP seems to have a per-TIC average solution time with linear dependence (R^2^≥0.89) on size of model and/or database used, see Figure 4(A) through (C). The same procedure was applied to the aggregated CPs’ solution time, but with significantly different results. Exponential trend lines were able to fit with a high correlation coefficient (R^2^≥0.96) between model, database, total system size, and CPs aggregated solution time. This is indicative of a strong correlation between CPs aggregate solution time number of reactions in the total system, and that increasing total system reactions greatly increases CPs aggregate solution time.

## DISCUSSION

Introduced here is a novel optimization-based tool, OptFill, which can be used to increase the automation of the curation of GSMs. This takes the form of both the whole OptFill tool, to automate the filling of metabolic gaps in a reconstructed model, and the mTFP, which can automate the identification of TICs for manual resolution. In this work, the OptFill tool was applied in sequence to three test models of increasing size as well as to a GSM of *E. coli*, *i*JR904. These applications highlighted the utility of OptFill, some solutions for holistically gapfilling metabolic models, the computational expense of the tool, and a method for reducing that expense.

This method has considerable potential to be adapted to other metabolic systems (eukaryotic and prokaryotic system) and is not specific to any identifier system such as KEGG or ModelSeed. For instance, while all test models as well as *i*JR904/*i*AF1260 have been prokaryotic systems, there is no reason why this approach would not similarly work in a eukaryotic organism. Further, the Supplemental Files, as well as the mathematical methods, are flexible enough that any system of reaction and metabolite identifiers, such as KEGG (Kanehisa *et al*., 2017), MetaCyc (Caspi *et al*., 2014), BIGG (King *et al*., 2016), K-Base (Arkin *et al*., 2018), or custom identifiers, may be used for metabolites and/or reactions, making this tool applicable to a wide variety of existing GSM-building methods. This has been demonstrated in that KEGG identifiers were used in the test models, whereas BIGG identifiers were used by the *i*JR904 and *i*AF1260 models (Reed *et al*., 2003)(Feist *et al*., 2007b).

From the observation of TFP solution times, it is evident that the TFP and mTFP could scale-up to genome-scale models of metabolism as a linear trend line strongly (R^2^≥0.89) describes the per-TIC solution time given the computational resources at hand. So long as the number of TICs in the system remains reasonable, this portion of OptFill is transferrable to large-scale GSM systems, or to situations where computational resources are limited. The transferability of the OptFill method as a whole is likely limited by the computational resources available to the end-user, as the aggregate solution time of the three CPs is well-described by an exponential trend line (R^2^≥0.97), suggesting that those without access to powerful computational resources may have difficulty implementing OptFill in a reasonable timeframe, unless, for instance the end-user makes trade-offs between the solution order (e.g. each subsequent solution is truly globally optimal) and solution time. These trade-off issues, such as shown in a minor way with the OptFilling of *i*JR904, may likely be fixed by more advanced MILP solvers which are currently available or by advances in optimization which may be made in future.

When implementing OptFill in other systems, a high quality model and database should be used for reasons of both limiting the number of solutions and limiting the time the OptFill method takes to complete. The primary problem is the number of feasible and unique combinations possible. For instance, if a multi-step reaction is included in a database in addition to its component reaction steps, this can potentially double the number of solutions found by both the TFP and CPs. To explain, if the multi-step reaction participates in *n* TICs, then its component step reactions would participate in *n* TICs. This results in *2n* TICs, where only *n* TICs need be identified. The same argument applies for CPs’ solutions. This error in model reconstruction could then double (or more) the number of TICs and CPs’ solutions as well as total OptFill runtime in a stroke. In larger models, such issues can result in a significant expenditure of time, potentially days, and computational resources which need not be expended should the model and database used be of high quality. This is shown in this work in the failure to achieve a reasonable number of solutions or reasonable solution times in the OptFilling of *i*JR904 with a poorly curated database based on *i*AF1260; however, when the database was better curated, reasonable numbers of solutions and solution times were achieved. Therefore, it is important to address as many inherent TICs which occur both in the model and in the database as is feasible using the mTFP on both model and database to identify and address these TICs.

While throughout this text reaction cycling in the absence of nutrition, that is Thermodynamically Infeasible Cycling, is described as and assumed to be a phenomena which is to be avoided in GSMs, this is not always the case. In many biological systems, cycling of some type does occur, and the lack of that cycling may decrease the accuracy of a metabolic model. However, cycles included in a GSM should be carefully considered with respect to their biological relevance, their magnitude, and their effect, particularly when they occur in the absence of nutrition provided to the model. In essence, this work can be used to remove and/or avoid all cycling which can occur in the absence of nutrition provided to the model, or can be used to ensure that cycles retained are deliberate and have biological relevance if included. If cycles occur in a GSM model in the absence of nutrition provided to the model and are biologically relevant, best practice should be to use other literature data available to limit the scope of the cycling to feasible number, as the default bounds on most reactions in a GSM generally represent infeasibly large reaction rates.

In future, this work will be used as a gapfilling and curation strategy for the development of GSMs. In concert with advances in optimization solvers and available computational power, these methods, the TFP, CPs, and their modified versions, will provide an alternative holistic method of model curation. At present, those model-building tools with high computational power at their disposal, such as ModelSeed (Overbeek *et al*., 2005) and K-Base (Arkin *et al*., 2018), may well be able to implement OptFill and its components for large GSMs to improve their automated curation capabilities.

## ACKNOWLEDGEMENTS

This work has been completed utilizing the Holland Computing Center of the University of Nebraska, which receives support from the Nebraska Research Initiative. The authors gratefully acknowledge funding from UNL Faculty Startup Grant 21-1106-4038.

## AUTHOR CONTRIBUTIONS

Experiments have been conceived by R.S. and W.L.S. W.L.S. designed the problems, performed the experiments, and analyzed the data. R.S. and W.L.S. wrote the manuscript.

## MAIN TABLES AND LEGENDS

This manuscript contains no tables.

## STAR METHODS

### KEY RESOURCES TABLE

| REAGENT or RESOURCE | SOURCE | IDENTIFIER |
| --- | --- | --- |
| Software and Algorithms |  |  |
| Generalized Algebraic Modeling System (GAMS) version 24.7.4 | GAMS Development Corp. (URL: www.gams.com/) | N/A |
| CPLEX Solver version 12.6 | GAMS Development Corp. (URL: www.gams.com/) | N/A |
| Perl version 5.26 (for Unix) | Perl.org (URL: www.perl.org) | N/A |
| Strawberry Perl version 5.24.0.1 (for Windows) | Perl.org (URL: http://strawberryperl.com/) | N/A |
| The world-wide-web library for Perl, module 6.39 | Perl meta::cpan (URL: metacpan.org/pod/LWP) | N/A |
| Python version 3.3 (for Unix) | Python Software Foundation (URL: www.python.org) | N/A |
| Kyoto Encyclopedia of Genes and Genomes (KEGG) | Kanehisa Laboratories (URL: www.kegg.jp) | RRID:SCR_012773 |
| iJR904 | Reed et al. 2003 | N/A |
| Other |  |  |
| Holland Computing Center, Crane Cluster (64 GB RAM, Intel Xenon E5-2670 2.60 GHz processor, 2 CPUs per node) | Holland Computing Center, University of Nebraska (URL: hcc.unl.edu/) | N/A |

### LEAD CONTACT AND MATERIALS

Further information and requests for resources and reagents should be directed to and will be fulfilled by the Lead Contact, Rajib Saha.

#### Materials Availability Statement

This study has produced several unique software codes in the form of GAMS, Perl, or Python programming languages/tools. These are included in the Supplemental Information. Further, the models and databases used in this study are also included in Supplemental Information.

### METHOD DETAILS

#### Model-Database TIC-Finding Problem (TFP)

The first step of the OptFill method requires the iterative solving of the Mixed Integer Linear Programming (MILP) TIC-Finding Problem (TFP) applied to the model and database. This problem is defined below and is designed such that a TIC which could exist between the model and database with given reaction flux bounds will be a solution to the TFP.

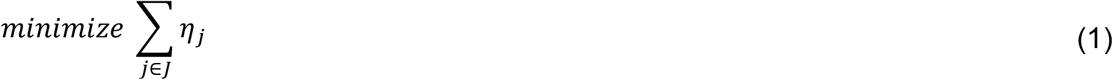

Subject to (s.t.)

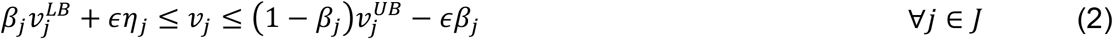

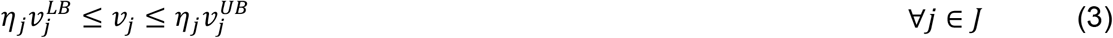

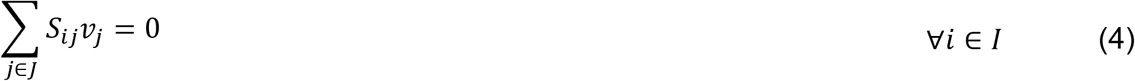

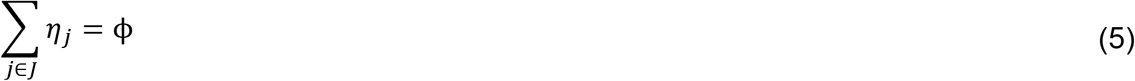

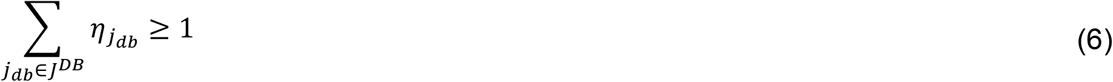

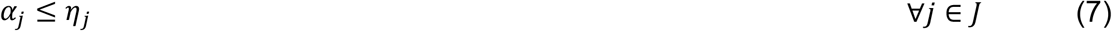

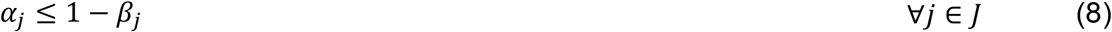

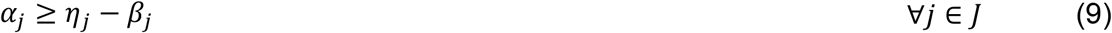

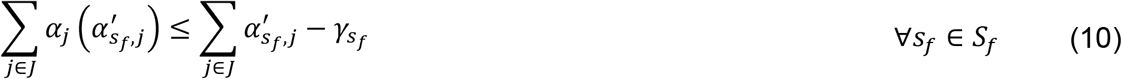

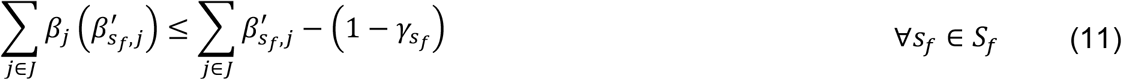

Fixed Values

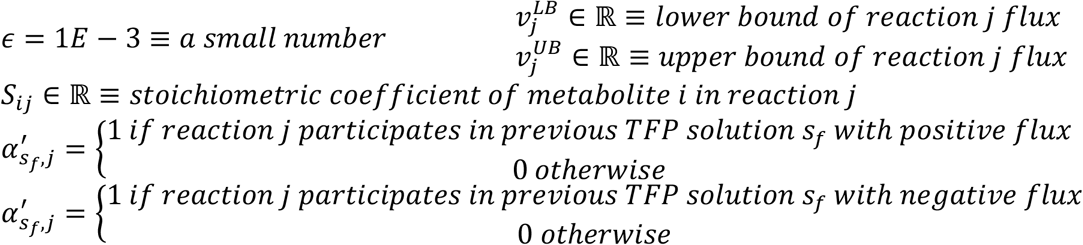

Variables

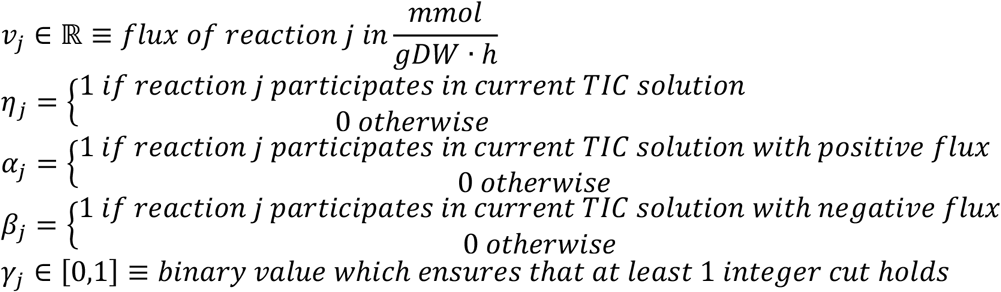

The set *s_f_* is the set of all previously found TICs and represents the solution space that is known. It should be noted that set *J* is the set of all reactions in the database and model, of which set *J^DB^*, the set of all reactions in the database, is a subset. Further, it should be noted that *I* is the set of all metabolites in the database and the model. Parameters (fixed values) and variables are defined after all constraints have been listed. The TIC-finding problem is run with all nutrient uptakes turned off, so that any reaction flux is unrealistic and due to one or more TICs. The TFP is included in Supplemental File 4 as GAMS (Generalized Algebraic Modeling System) code. The following subsections will describe the above equations constituting the TFP in detail.

#### Objective function and sought TIC size

The solution of the TFP is itself a TIC. The objective function, equation (1), is minimization of the number of reactions participating in the TIC solution. This objective function is irrelevant in the solution due to equation (5), as equation (5) specifies the size of the TIC sought, and thus the objective function value, and is included to ensure that each possible TIC size is investigated. The order of solutions, when the workflow in Figure 2 is followed, is unimportant, and may vary each time the TFP is applied to a model.

#### Enforcing flux bounds and reaction participation

Equations (2) and (3) are constraints which enforce the given reaction flux bounds and determine if a reaction participates in the identified TIC. The variable *η_j_* stores if a reaction participates in a TIC, while variables *α_j_* and *β_j_* store direction of participation. Reaction flux bounds 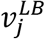 and 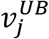 are determined manually based on reaction direction (reversible, irreversible forward, or irreversible backward), limitations on nutrient uptake rates, and reaction state (either on or off depending on genotype, nutrient availability). Equation (6) ensures that at least one database reaction holds flux. Equation (2) specifically identifies if reaction *j* participates in the solution TIC by requiring some small, minimum reaction flux, *ϵ*, for participating reactions such that equation (12) is true. Further, it identifies the direction of that reaction.

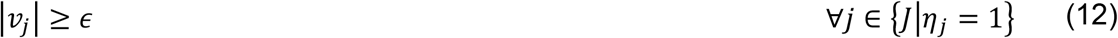

Equation (3) ensures that if any reaction does not meet the reaction flux threshold to participate in the TIC solution, that the reaction flux is constraint to zero.

#### Identifying positive flux participation in the TIC

Equations (7) through (9) are a linearized version of the following statement.

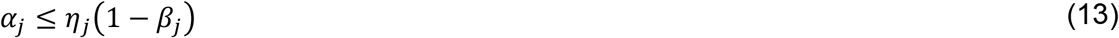

The linearization in equations (7) through (9) functions the same as (13) because *η_j_* and *β_j_* are binary variables. This linearization is made in order to preserve the linear nature of the TFP. A linear optimization problem can guarantee both global solution optimality and that all solutions in the solution space can be enumerated, which in this case guarantees that all TICs are found of a given size.

#### Integer Cuts for Repeated Solutions

Equations (10) and (11) are integer cuts which prevent repetition of solutions. It should be noted that these repeated solutions include direction. Therefore, to be identified as the same TIC, the set of participating reactions and the directions in which they participate must be the same. Consider the following set of chemical equations for an illustration of how these integer cuts prevent repeated solutions.

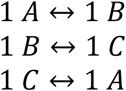

The TFP, because of these integer cuts, would identify two TICs existing in this set of chemical reactions. The first would be all reactions listed above proceeding in the forward direction, while the second would be all reactions listed above proceeding in the backward direction. These are identified separately because their reaction directions are different, although the participating reactions are the same.

#### Modified TIC-Finding Problem (mTFP)

The TFP can be modified for the identification of TICs inherent to a metabolic model to aid in model curation. The modified TIC-Finding Problem (mTFP) can be formulated via equations (1) through (5) and equations (7) through (11). All set, parameter, and variable definitions are the same as in the unmodified TFP.

#### First Connecting Problem (CP1)

The connecting problems are the series of optimization problems which are solved following the solving of the TFP. First discussed will be the first Connecting Problem (CP1). The solution to a CP is a set of database reactions which, when added to the model, will increase model connectivity. The solution to CP1 gives the maximum number of model metabolites which could be connected using the database. The formulation of CP1 is given below.

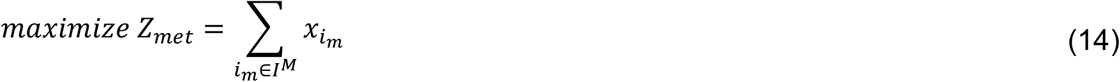

Subject to

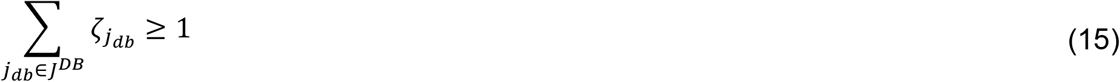

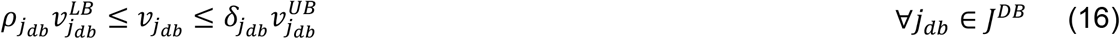

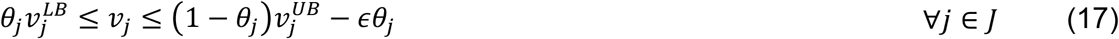

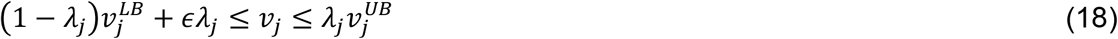

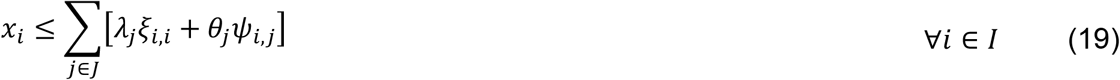

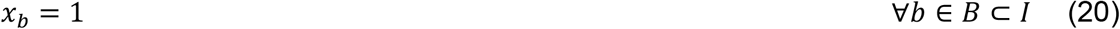

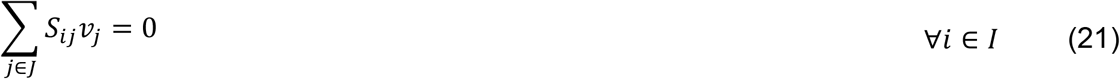

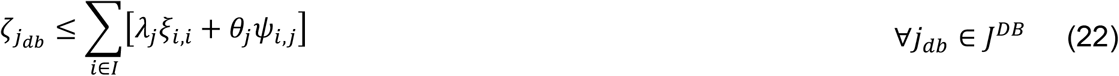

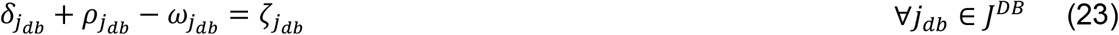

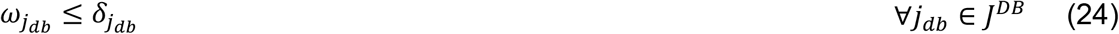

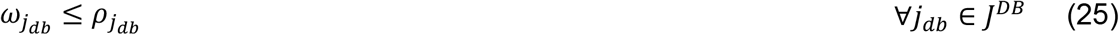

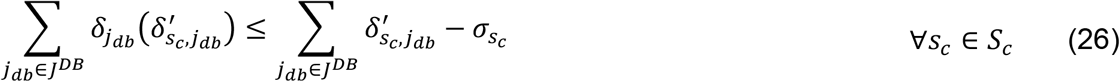

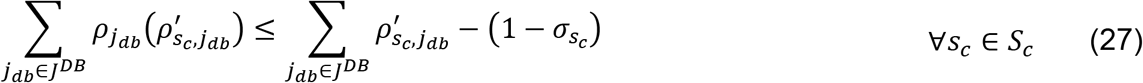

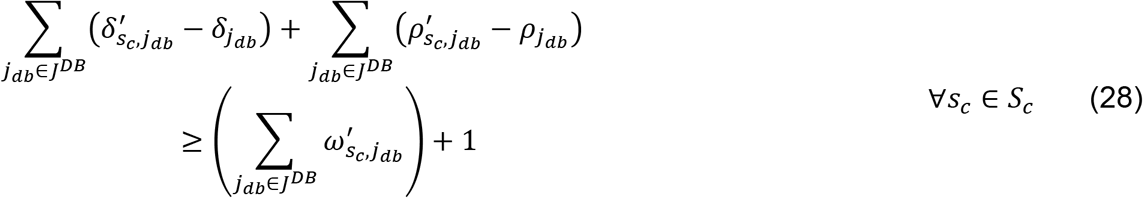

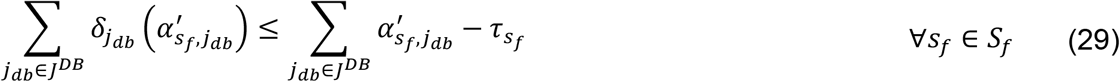

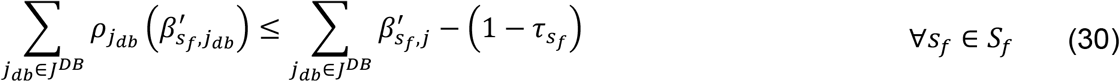

*Fixed Values Unique to CP1*

*M* = 1*E*3 ≡ *a very large number*

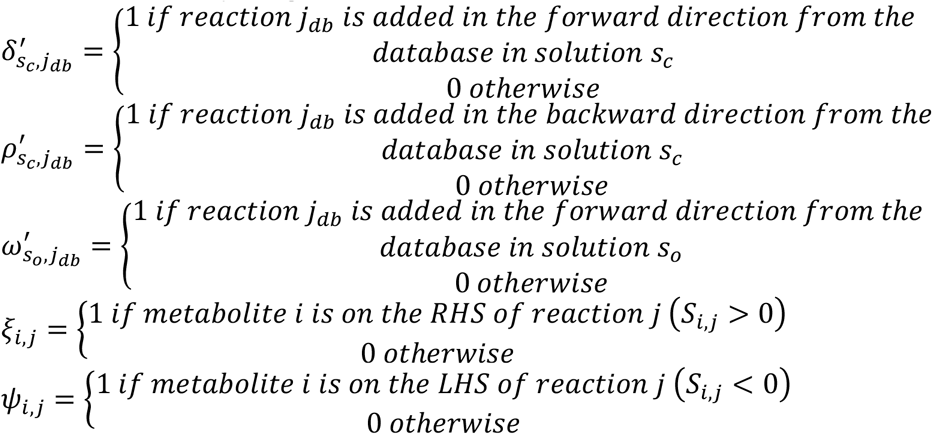

*Variables Unique to CP1*

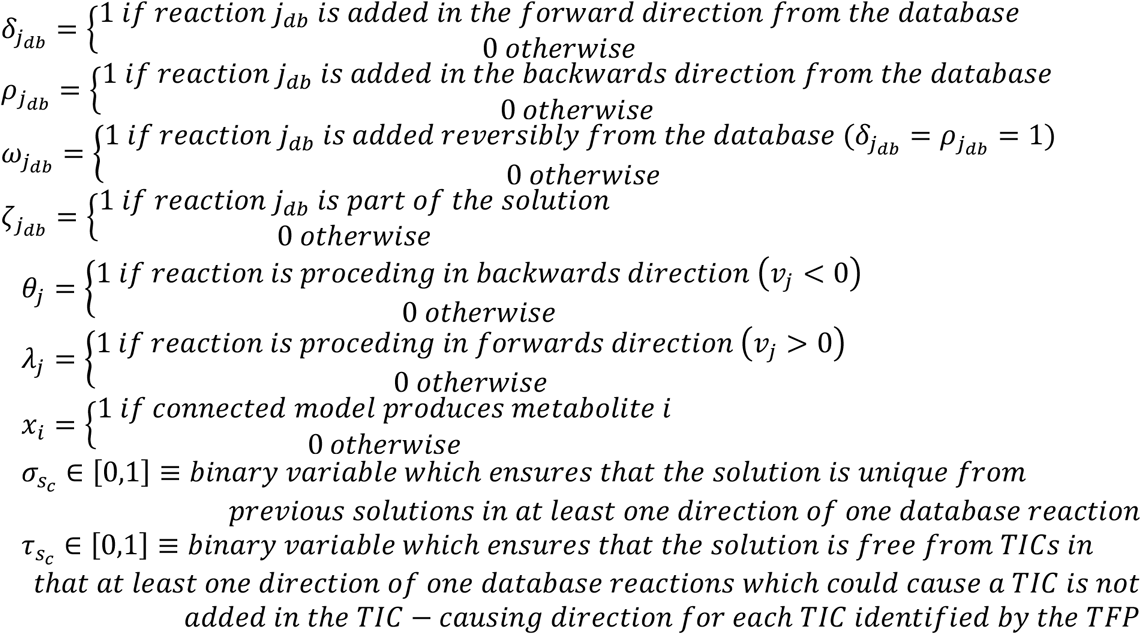

Where *I^M^* is defined as the set of metabolites in the model and is a subset of *I*. When CP1 is solved, the optimal value of *Z_met_* is the maximum number of metabolites which can be connected in the model by adding reactions from the database, given all previous solutions (if any) and all identified potential TICs. It should be noted that all sets and parameters have the same definitions here as in the TFP, with the additions of *J^M^* being the set of model reactions which is a subset of *J*, of *I^M^* being the set of model metabolites which is a subset of *I*, *s_c_* being the set of all previous connecting problem solutions, *s_o_* being the set of all previous connecting problem solutions with at least one reversible reaction being added from the database, and *B* being the set of all metabolites which are involved in the biomass equation which is a subset of *I*.

The following statements give, broadly, the rational for each constraint equation. Equation (14) ensures that at least one reaction is added from the database for each solution. Equation (15) ensures that each database reaction only has flux if it is added. Equation (16) ensures that the user-defined reaction flux bounds hold. Equations (17) through (19) determine which metabolites the fixed model can produce, equation (19) ensures that the fixed model can produce biomass. Equation (20) ensures mass balance. Equation (21) ensures that added reactions are productive, e.g. that the added reaction does produce one or more metabolites. Equations (22) through (24) ensure that each database reaction for the connecting solution is added as a forward, backward, or reversible reaction (e.g. both as a forward and a backward reaction). Equations (25) though (28) are integer cuts preventing repeated solutions, while Equations (29) and (30) are integer cuts preventing the full addition of a TIC through the CP solution. The following subsections will describe some of the above equations constituting the CPs in greater detail. The CPs are included in Supplemental File 4 as GAMS (Generalized Algebraic Modeling System) code. The following subsections will describe some of the above equations constituting the CPs in greater detail.

#### Determination of Metabolite Production

Important to CPs is the determination of whether or not a metabolite is produced in the connected model. Equations (17) and (18) are used to determine which direction reactions proceed in the connected model. Equation (19) essentially states that a metabolite is produced if at least one reaction produces that metabolite by having flux in the direction of that metabolite (either through backwards flux and a negative stoichiometric coefficient or forward flux and a positive stoichiometric coefficient). Equation (20) ensures that all metabolites necessary for growth (those involved in biomass production) are produced, as all models of metabolism should be capable of producing biomass, even if biomass is not ultimately the objective used. For instance, alternate objectives could include the maximization of production of a given metabolite (Herrgård, Fong and Palsson, 2006)(Price, Reed and Palsson, 2004), the minimization of the uptake of a particular substrate (Gomes de Oliveira Dal’Molin *et al*., 2015), or minimization of metabolic adjustment (MOMA) (Herrgård, Fong and Palsson, 2006)(Price, Reed and Palsson, 2004). Ultimately, each objective type some fixed or variable non-zero level of biomass production and therefore all models require some ability to grow, making these constraints reasonable for reconstructions regardless of the ultimate objective used. Equation (22) ensures that reactions added from the database are productive, e.g. that each added reaction is capable of producing at least one metabolite. This constraint ensures that reactions incapable of carrying flux are not added to the model.

#### Direction of Added Database Reactions

Equations (22) through (25) largely deal with the direction in which reactions are added from the database. Equations (22) ensures that reactions added from the database are productive. Equation (23) ensures that *ζ_jdb_* is equal to 1 if reaction *j_db_* is added to the model as part of this solution, and zero otherwise. Equations (23) through (25) are the linearization of the multiplication of two binary variables stated below.

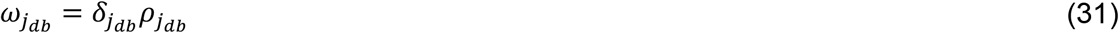

This linearization is done for the same reasons that the TFP has been linearized. The sum of these constraints ensures that any reaction added reversibly is treated as a reaction added both forward and backwards for the purposes of integer cuts to avoid repeated solutions.

#### Integer Cuts for Repeated Solutions

Equations (26) through (28) define integer cuts used to avoid repeat solutions. Equations (26) and (27) have been designed on similar lines to (10) and (11), designed to avoid repeat solutions. Through the integer cuts in equations (26) through (28), both the reactions and their directions are integral to the solution; therefore, any different between solutions in reaction direction or reactions included is recorded as a second solution. Equation (28) prevents the repetition of a solution that could be caused by changing a reversible database reaction addition into an irreversible one.

#### Integer Cuts for TIC-less Connecting

Equations (29) through (30) define integer cuts which ensure that a TIC is not added to the connecting solution. This is done by considering both reaction identity and direction for both the addition of database reactions and for the avoidance of TICs. This results in a minimum perturbation to the solution space of CPs caused by each TIC. As with other directional integer cuts, only one cut needs be in effect at minimum in order to define a new solution.

#### Modified First Connecting Problem

A modified CP1 was used to get an initial count of the maximum number of metabolites which the raw model can produce. This modified CP1 made use of equations (14), and (16) through (30). In place of equation (15) the following equation was used to ensure that no database reactions were considered in maximizing the number of metabolites which may be connected.

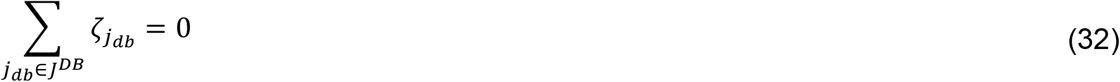

#### Second Connecting Problem

The second Connecting Problem (CP2) is defined as equations (15) through (30) with the addition of the objective function and constraint equation (34) stated below.

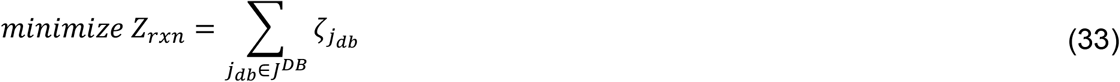

s.t.

Equations (15) through (30)

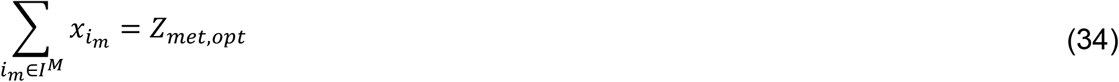

Where *Z_met,opt_* is defined as the optimal objective value of CP1. When CP2 is solved, the optimal value of *Z_rxn_* is the minimum number of reactions which, when added from the database, can connect the previously determined maximum number of model metabolites, given all previous solutions (if any) and all identified potential TICs.

#### Third Connecting Problem

The third Connecting Problem (CP3) is defined as equations (15) through (30), equation (34), and constraint equation (36) stated below.

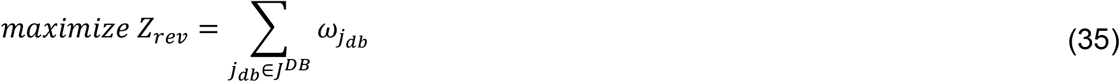

s.t.

Equations (15) through (31), (34)

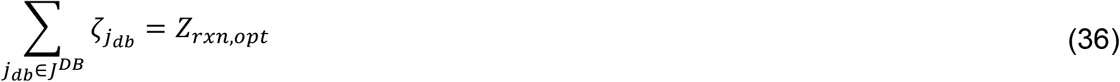

Where *Z_rxn,opt_* is defined as the optimal objective value of CP2. When CP3 is solved, the optimal value of *Z_rev_* is the maximum number of reversible reactions which can be used to achieve the minimum number of reaction additions to maximize model connectivity, given all previous solutions (if any) and all identified potential TICs. The solution of CP3 is the solution accepted as optimal.

CP3 has been found to be needed due to allowing database reactions to be added forward, backward, and reversibly. Since adding a reaction reversibly rather than irreversibly in some cases has made no difference, this resulted in an inconsistent number of solutions to the set of CPs. Therefore, in one run two solutions would be returned (the irreversible solution has been returned, then the reversible), where in a subsequent run perhaps only one solution would be returned if the reversible solution has been returned first, and then integer cuts (26) and (27) would preclude the irreversible solution. This third connecting problem has been added to deal with such situations by forcing the reversible solution to be returned first, resulting in a standardized, minimized set of solutions.

#### FBA of Connected Model

Once the CPs have been solved and the identity and direction of models to be added from the database to the model for a given solution are known, Flux Balance Analysis (FBA) is performed on the connected model. As the models are not physically merged, this takes the following form.

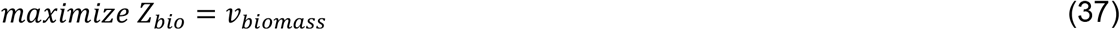

s.t.

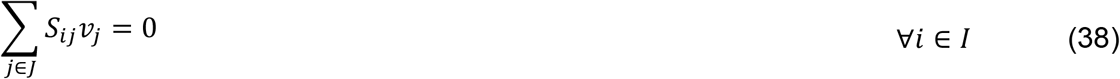

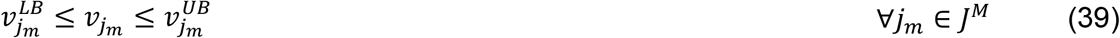

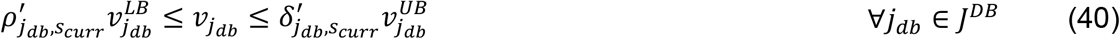

All variables, parameters, and sets are the same as in previous equations, and in addition *s_curr_* represents the current connecting solution. In the above formulation, equation (39) takes into account the current solution of the CPs. A biomass maximization objective function was chosen for this work, but other objective could be selected depending on what part of metabolism is of most interest.

#### Creation of Test Models and Databases

Test model have been created in tandem with their databases using KEGG maps of pathways to identify sets of reactions which might produce a functional metabolic model. The first Test Model (TM1) and Test Database (TDb1) have been built from the “starch and sucrose metabolism” (map00500) and the “glycolysis/gluconeogenesis” (map00010) metabolic maps with the goal of producing a minimal prokaryotic model which utilizes sucrose, produces ethanol and biomass, and has some TICs which exist between the database and model where TM1 cannot produce biomass (without some TDb1 reactions) and contains no inherent TICs. Since only sucrose metabolism and glycolysis have been included in this model, biomass for this model is based on glucose, fructose, and an arbitrary growth-associated maintenance (GAM) value of 2. The coefficient of glucose in the biomass equation has then been scaled such that the molecular weight of biomass is 1000 g/mol. Non-Growth Associated Maintenance (NGAM) has also been defined arbitrarily as 2. TM1 and TDb1 have been constructed rationally with as many reversible reactions as possible, such that 22 of the 28 reactions are reversible in TM1 and all 17 reactions are reversible in TDb1. Once TM1 and TDb1 have been constructed, OptFill has been applied to them. This has resulted in the identification of 31 TICs consisting of 3 to 12 reactions by the TFP using the CPLEX solver. See results section for detail.

The first solution reported by OptFill for TM1/TDb1 has been added to TM1 to create the initial second Test Model (TM2). Added manually to this initial TM2 model is portions of the “pentose phosphate” pathway (map00030) and fatty acid biosynthesis” (map00061) pathway. The biomass equation has been updated to include a small amount (stoichiometric coefficient 0.01) of each new fatty acid product (8-, 10-, 12-, 14-, 16-, and 18-carbon fatty acid products) and the coefficient of glucose has again been adjusted to ensure biomass molecular weight was 1000 g/mol. Certain reactions in both pathways have been selected to constitute the second Test Database (TDb2), again with the aim of being a small prokaryotic model which utilizes sucrose, produces ethanol, produces biomass, and has some TICs which exist between the database and model where TM2 cannot produce biomass (without some TDb2 reactions) and contains no inherent TICs. In total, TM2 consists of 77 reactions (with 65 being reversible), and TDd2 consists of 34 reactions (all reversible). Once TM2 and TDb2 have been constructed, OptFill has been applied to them, see results section for details.

As with the construction of TM2, the third Test Model (TM3) has initially been constructed from the first solution of OptFill applied to TM2/TDb2 added to a test model. This test model has then been expanded to include “nitrogen metabolism” (map00910, with ammonium uptake), “sulfur metabolism” (map00920, with sulfate uptake), and synthesis pathways for all 20 amino acids. The biomass equation has been updated to include a small amount (stoichiometric coefficient 0.1) of each of the 20 primary amino acids, following which the coefficient of glucose has again been adjusted to ensure biomass molecular weight was 1000 g/mol. Unlike previous test models, this working test model (e.g. capable of producing biomass) with some TICs has first been developed, split between old reactions (belonging to TM2 or OptFill solution thereof) and new reactions. Then each new reaction has been assigned a random value (between 0 and 1) and those with a value greater than or equal to 0.7 have been assigned to the third Test Database (TDb3), and those with a value less than or equal to 0.8 have been assigned to the third Test Model (TM3). The code to perform this is included as part of Supplemental File 3. Following this, the mTFP has been applied to TM3 in order to ensure that the model is TIC-less. For removing TICs from TM3, the number of occurrences of each reaction participating in all TICs has been counted, that has the highest occurrence, excluding those reactions from TM2 and TDb2, has been moved to TDb3. In the case of ties, the reaction with the highest reaction ID number has been moved to TDb3. In total, TM3 consists of 210 reactions (196 reversible), and TDb3 consists of 77 reactions (all reversible). Once TM3 and TDb3 have been constructed, OptFill has been applied to them, see results section for details.

It should be noted that for all instances of OptFill applied to test models some low number of execution errors have been allowed, five are allowed in this example option allowing execution errors: “execerr=5”. This has been done because GAMS throws an execution error if the RHS and LHS of a constraint are fixed and those fixed values do not satisfy the constraint. In the case of OptFill, this is not necessarily an issue, as it simply indicates that there are no more feasible solutions and that the program should continue onto the next problem or step. Graphical summaries comparing project runtimes have then been generated in Supplemental File 5 (Microsoft Excel) to produce Figure 4. Trend line and Pearson correlation values included in this figure have been generated automatically by Microsoft Excel. Linear, logarithmic, exponential, and power trend lines have been investigated, and the best fit line is displayed for each dataset. Polynomial trend lines have not been investigated as these trend lines can lead to overfitting errors.

#### Application of *i*AF1260 to *i*JR904

In the application of OptFill to published *Escherichia coli* GSMs, *i*JR904 (Reed *et al*., 2003) was treated as the model and *i*AF1260 (Feist *et al*., 2007a) as the source of reactions to build the database for OptFill. Minor formatting of both of these models was accomplished using the code in Supplemental File 1. Such formatting changes include changing of how reaction arrows appeared and location of metabolite compartment notation. Following this formatting, all exchange reactions were removed from *i*AF1260, as it was decided to use the media definition provided for *i*JR904 by Reed *et al*., 2003, specifically for the case of aerobic growth on acetate. Whereas very large bounds in *i*JR904 have been defined as 1e^30^, these have been redefined as 1e^3^ as both quantities are sufficiently large in the context of GSMs to be a red flag should any reaction flux reach that quantity. Further, 1e^3^ is the value of *M* used elsewhere in the code, resulting in a standard value for a “very large number”.

Once the aforementioned changes had been made, *i*AF1260 (sans exchange reactions) and *i*JR904 were compared in Supplemental File 5 so that reactions that are in both model would be removed from *i*AF1260. These modifications resulted in 1441 reactions remaining in the initial *i*AF1260-based database. The initial *i*AF1260-based database is provided in Supplemental File 3, as is the GAMS code used in this application of OptFill. The OptFilling of *i*JR904 using an *i*AF1260-based database is different from the code used for the test models/database only in formatting of the output file (identifiers used were considerably longer than KEGG identifiers causing formatting issues). This was allowed for seven days to attempt to solve, in which time it did not return a single CPs solution; therefore, it was decided that the database needed to be made smaller. Both the initial *i*AF1260-based database and *i*JR904 were combined into a single pseudo-model file, to which Flux Variability Analysis (FVA) was applied. Those reactions which hold flux, 715 reactions, formed the second *i*AF1260-based database.

OptFill was applied to this second database, but still resulted in very long solution times; therefore, those reactions which participated in solutions which were achieved in 24 hours (four solutions) were chosen to form the final *i*AF1260-based database. This database consists of 182 reactions. It was found that this resulted in no TFP solutions; therefore, six more reactions were added to produce a database which had 25 potential TICs with the *i*JR904 model. OptFill was then applied to *i*JR904 using this final *i*AF1260-based database of 188 reactions.

#### CPLEX Solver Options

As the order of solutions presented is important, solver options which allowed non-optimal solutions or created relaxations by which the truly optimal solution could not be reached, or a sub-optimal solution would be accepted, were disabled. In particular, the infeasibility gap was set to the lowest possible value, small infeasibilities were disallowed, no relaxation was allowed in the value of integers, no optimality gap was allowed in the solution, and solver cuts which could result in non-optimal solutions were disabled. These cuts included zero-half, flow, clique, cover, mixed integer rounding, GUB cover, and Gomory fractional cuts. While the lack of these relaxation options and cuts no doubt increased solution time, these relaxations would decrease solution accuracy and order which was deemed unacceptable. The list of CPLEX relaxations used in this work can be found in Supplemental File 3.

#### Available Hard- and Soft-ware Tools

The University of Nebraska-Lincoln Holland Computing Center Crane Cluster was used. The nodes used on this cluster for this work have 64 GB RAM, Intel Xenon E5-2670 2.60 GHz processors, and 2 CPUs per 16 nodes for 548 nodes. In addition, Crane uses an old version of GAMS, version 24.7.4 released in March 2016 as opposed to the current version 28.1.0 released August 2019; therefore, algorithms used in this work may be sub-optimal compared to those which may be available at present. Further, this version of GAMS corresponds to CPLEX library 12.6, which was release in January 2014. Therefore, solution time may be different, and significantly so, for users with access to different hard- and soft-ware tools. It should also be noted that the solution times are not static. That is, each time OptFill is run, it may have different values for solution times; however, the patterns of solution time and distributions should remain relatively consistent.

## Acronyms Used

LHS: Left Hand Side
RHS: Right Hand Side
TFP: TIC-Finding Problem
CPs: Connecting Problems
FBA: Flux Balance Analysis
TM1: First Test Model
TDb1: First Test Database
TM2: Second Test Model
TDb2: First Test Database
TM3: Third Test Model
TDb3: First Test Database
GAM: Growth Associated Maintenance
NGAM: Non-Growth Associated Maintenance

## QUANTIFICATION AND STATISTICAL ANALYSIS

The software used to calculate statistics presented in this work was GAMS; however, as GAMS has no built-in statistics tools, these calculations were performed in the code included in Supplemental File 4. The two statistics calculated include mean and standard deviation. In this study, the arithmetic mean was used, and the standard deviation calculation used was that of a population (as opposed to a sample), as the full population of solutions was used in the test statistic.

## DATA AND CODE AVAILABILITY

All code used as part of this work has been included in the Supplemental Information of this work. Due to limitations of the number of supplemental materials allowed, some code and input data of this work has been combined in such a way that they will need to be separated before replicating this work. Supplemental File 6 has information on how to accomplish this process.

## SUPPLEMENTAL VIDEO, DATA, AND EXCEL TABLE TITLE AND LEGENDS

**Supplemental File 1: Perl code for formatting of iJR904 and iAF1260**. Some formatting was required of iJR904 and iAF1260 to put them in a form readable by current OptFill code. This piece of Perl code accomplishes this task.

**Supplemental File 2: iJR904**. This supplemental file is built from the supplemental file provided by Reed et al. 2003 with the addition of columns defining metabolite IDs used by Reed et al. 2003 with KEGG identifiers. This substitution was done in this study.

**Supplemental File 3: Zip folder of other important files**. Due to restriction on the number of allowable supplemental files, this single zip file contains the content of what is many individual files, including code, solver options, and test-file results. Consult Supplemental File 6 for what all is contained within Supplemental File 3 and the function or purpose of each item.

**Supplemental File 4: GAMS Version of OptFill**. This file is the executable GAMS code for the application of OptFill to a given model. Before execution, consult Supplemental File 6 for how to rebuild all files necessary for this code to properly execute.

**Supplemental File 5: Result summaries, graphs and biomass calculations**. Microsoft Excel workbook which was used to calculate biomass composition for the test models as well as generate graphs used in Figures A and B.

**Supplemental File 6: Recreating the digital environment necessary to replicate this study**. This file is a Microsoft Word document which discusses how to create each of the models and databases used in this study from Supplemental File 3. This process was considered too tedious to include in the main text, but is necessary for replication.

## REFERENCES

Andersen, M. R., Nielsen, M. L. and Nielsen, J. (2008) ‘Metabolic model integration of the bibliome, genome, metabolome and reactome of Aspergillus niger’, Molecular Systems Biology, 4(178), p. 178. doi: 10.1038/msb.2008.12.

Arkin, A. P. et al. (2018) ‘KBase: The United States department of energy systems biology knowledgebase’, Nature Biotechnology, 36(7), pp. 566–569. doi:10.1038/nbt.4163.

Beyer, P. et al. (2002) ‘Golden Rice: introducing the beta-carotene biosynthesis pathway into rice endosperm by genetic engineering to defeat vitamin A deficiency.’, The Journal of nutrition, 132(3), pp. 506S–510S. Available at: http://www.ncbi.nlm.nih.gov/pubmed/11880581.

Brunk, E. et al. (2018) ‘Recon3D enables a three-dimensional view of gene variation in human metabolism’, Nature Biotechnology. Nature Publishing Group, 36(3), pp. 272–281. doi: 10.1038/nbt.4072.

Burgard, A. P., Pharkya, P. and Maranas, C. D. (2003) ‘OptKnock: A Bilevel Programming Framework for Identifying Gene Knockout Strategies for Microbial Strain Optimization’, Biotechnology and Bioengineering, 84(6), pp. 647–657. doi: 10.1002/bit.10803.

Caspi, R. (2006) ‘MetaCyc: a multiorganism database of metabolic pathways and enzymes’, Nucleic Acids Research, 34(90001), pp. D511–D516. doi: 10.1093/nar/gkj128.

Caspi, R. et al. (2014) ‘The MetaCyc database of metabolic pathways and enzymes and the BioCyc collection of Pathway/Genome Databases’, Nucleic Acids Research, 42(D1), pp. 459–471. doi: 10.1093/nar/gkt1103.

Chowdhury, R., Chowdhury, A. and Maranas, C. D. (2015) ‘Using gene essentiality and synthetic lethality information to correct yeast and CHO cell genome-scale models’, Metabolites, 5(4), pp. 536–570. doi: 10.3390/metabo5040536.

Cuevas, D. A. et al. (2019) ‘Elucidating genomic gaps using phenotypic profiles [version 2; peer review: 1 approved, 1 approved with reservations]’, (May), pp. 1–28.

Feist, A. M. et al. (2007a) ‘A genome-scale metabolic reconstruction for Escherichia coli K-12 MG1655 that accounts for 1260 ORFs and thermodynamic information’, Molecular Systems Biology, 3(121), pp. 1–18. doi: 10.1038/msb4100155.

Feist, A. M. et al. (2007b) ‘A genome-scale metabolic reconstruction for Escherichia coli K-12 MG1655 that accounts for 1260 ORFs and thermodynamic information’, Molecular Systems Biology, 3(121), pp. 1–18. doi: 10.1038/msb4100155.

Gomes de Oliveira Dal’Molin, C. et al. (2015) ‘A multi-tissue genome-scale metabolic modeling framework for the analysis of whole plant systems’, Frontiers in Plant Science, 6(January), pp. 1–12. doi: 10.3389/fpls.2015.00004.

Gudmundsson, S., Agudo, L. and Nogales, J. (2017) Applications of genome-scale metabolic models of microalgae and cyanobacteria in biotechnology, Microalgae-Based Biofuels and Bioproducts: From Feedstock Cultivation to End-Products. Elsevier Ltd. doi: 10.1016/B978-0-08-101023-5.00004-2.

Gudmundsson, S. and Thiele, I. (2010) ‘Computationally efficient flux variability analysis’, BMC Bioinformatics, 11(2), pp. 2–4. doi: 10.1186/1471-2105-11-489.

Hall, R. D., Brouwer, I. D. and Fitzgerald, M. A. (2008) ‘Plant metabolomics and its potential application for human nutrition’, Physiologia Plantarum, 132(2), pp. 162–175. doi: 10.1111/j.1399-3054.2007.00989.x.

Henry, C. S. et al. (2010) ‘High-throughput generation, optimization and analysis of genome-scale metabolic models’, Nature Biotechnology. Nature Publishing Group, 28(9), pp. 977–982. doi: 10.1038/nbt.1672.

Herrgård, M. J., Fong, S. S. and Palsson, B. (2006) ‘Identification of genome-scale metabolic network models using experimentally measured flux profiles’, PLoS Computational Biology, 2(7), pp. 0676–0686. doi: 10.1371/journal.pcbi.0020072.

Kanehisa, M. et al. (2017) ‘KEGG: New perspectives on genomes, pathways, diseases and drugs’, Nucleic Acids Research, 45(D1), pp. D353–D361. doi: 10.1093/nar/gkw1092.

Karp, P. D., Weaver, D. and Latendresse, M. (2018) ‘How accurate is automated gap filling of metabolic models?’, BMC Systems Biology. BMC Systems Biology, 12(1), pp. 1–11. doi: 10.1186/s12918-018-0593-7.

Kim, T. Y. et al. (2012) ‘Recent advances in reconstruction and applications of genome-scale metabolic models’, Current Opinion in Biotechnology. Elsevier Ltd, 23(4), pp. 617–623. doi: 10.1016/j.copbio.2011.10.007.

King, Z. A. et al. (2016) ‘BiGG Models: A platform for integrating, standardizing and sharing genome-scale models’, Nucleic Acids Research, 44(D1), pp. D515–D522. doi: 10.1093/nar/gkv1049.

Latendresse, M. and Karp, P. D. (2018) ‘Evaluation of reaction gap-filling accuracy by randomization’, BMC Bioinformatics. BMC Bioinformatics, 19(1), pp. 1–13. doi: 10.1186/s12859-018-2050-4.

Limviphuvadh, V. et al. (2018) ‘Discovering novel SNPs that are correlated with patient outcome in a Singaporean cancer patient cohort treated with gemcitabine-based chemotherapy’, BMC Cancer. BMC Cancer, 18(1), pp. 1–16. doi: 10.1186/s12885-018-4471-x.

Liu, J. et al. (2013) ‘Genome-scale reconstruction and in silico analysis of Aspergillus terreus metabolism’, Molecular BioSystems, 9(7), p. 1939. doi: 10.1039/c3mb70090a.

Magnúsdóttir, S. et al. (2016) ‘Generation of genome-scale metabolic reconstructions for 773 members of the human gut microbiota’, Nature Publishing Group, (November 2016). doi: 10.1038/nbt.3703.

De Martino, D. et al. (2013) ‘Counting and correcting thermodynamically infeasible flux cycles in genome-scale metabolic networks’, Metabolites, 3(4), pp. 946–966. doi: 10.3390/metabo3040946.

Ng, C. Y. et al. (2012) ‘Production of 2,3-butanediol in Saccharomyces cerevisiae by in silico aided metabolic engineering’, Microbial Cell Factories, 11(1), p. 68. doi: 10.1186/1475-2859-11-68.

Nigam, R. and Liang, S. (2007) ‘Algorithm for perturbing thermodynamically infeasible metabolic networks’, Computers in Biology and Medicine, 37(2), pp. 126–133. doi: 10.1016/j.compbiomed.2006.01.002.

Orth, J. D., Thiele, I. and Palsson, B. O. (2010) ‘What is flux balance analysis?’, Nature Publishing Group. Nature Publishing Group, 28(3), pp. 245–248. doi: 10.1038/nbt.1614.

Overbeek, R. et al. (2005) ‘The subsystems approach to genome annotation and its use in the project to annotate 1000 genomes’, Nucleic Acids Research, 33(17), pp. 5691–5702. doi: 10.1093/nar/gki866.

Pitkänen, E. et al. (2014) ‘Comparative Genome-Scale Reconstruction of Gapless Metabolic Networks for Present and Ancestral Species’, PLoS Computational Biology, 10(2). doi: 10.1371/journal.pcbi.1003465.

Price, N. D., Reed, J. L. and Palsson, B. (2004) ‘Genome-scale models of microbial cells: Evaluating the consequences of constraints’, Nature Reviews Microbiology, 2(11), pp. 886–897. doi: 10.1038/nrmicro1023.

Ranganathan, S., Suthers, P. F. and Maranas, C. D. (2010) ‘OptForce: An optimization procedure for identifying all genetic manipulations leading to targeted overproductions’, PLoS Computational Biology, 6(4). doi: 10.1371/journal.pcbi.1000744.

Reed, J. L. et al. (2003) ‘An expanded genome-scale model of Escherichia coli K-12 (iJR904 GSM/GPR).’, Genome biology, 4(9), pp. 1–12.

Saha, R. et al. (2012) ‘Reconstruction and Comparison of the Metabolic Potential of Cyanobacteria Cyanothece sp. ATCC 51142 and Synechocystis sp. PCC 6803’, PLoS ONE, 7(10). doi: 10.1371/journal.pone.0048285.

Saha, R. et al. (2016) ‘Diurnal Regulation of Cellular Processes in the Cyanobacterium Synechocystis sp. Strain PCC 6803: Insights from Transcriptomic’, MBio, 7(3), pp. 1–14. doi: 10.1128/mBio.00464-16.Editor.

Saha, R., Suthers, P. F. and Maranas, C. D. (2011) ‘Zea mays iRS1563: A comprehensive genome-scale metabolic reconstruction of maize metabolism’, PLoS ONE, 6(7). doi: 10.1371/journal.pone.0021784.

Satish Kumar, V., Dasika, M. S. and Maranas, C. D. (2007) ‘Optimization based automated curation of metabolic reconstructions’, BMC Bioinformatics, 8, pp. 1–16. doi: 10.1186/1471-2105-8-212.

Schellenberger, J., Lewis, N. E. and Palsson, B. (2011) ‘Elimination of thermodynamically infeasible loops in steady-state metabolic models’, Biophysical Journal. Biophysical Society, 100(3), pp. 544–553. doi: 10.1016/j.bpj.2010.12.3707.

Shoaie, S. et al. (2013) ‘Understanding the interactions between bacteria in the human gut through metabolic modeling’, Scientific reports, 3(2532), pp. 1–10. doi: 10.1038/srep02532.

Simons, M. et al. (2014) ‘Assessing the Metabolic Impact of Nitrogen Availability Using a Compartmentalized Maize Leaf Genome-Scale Model’, Plant Physiology, 166(3), pp. 1659–1674. doi: 10.1104/pp.114.245787.

Srinivasan, S., Cluett, W. R. and Mahadevan, R. (2015) ‘Constructing kinetic models of metabolism at genome-scales: A review’, Biotechnology Journal, 1359, pp. 1345–1359. doi: 10.1002/biot.201400522.

Thiele, I. and Palsson, B. Ø. (2010) ‘A protocol for generating a high-quality genome-scale metabolic reconstruction.’, Nature protocols. Nature Publishing Group, 5(1), pp. 93–121. doi: 10.1038/nprot.2009.203.

UniProtKB (2018) E. coli K12. Available at: www.uniprot.org/uniprot/?query=E.+coli+K-12+strain+1655&sort=score (Accessed: 20 August 2008).

