## Supplemental File 1 for "A Novel Optimization-Based Tool to Automate Infeasible Cycle-Free Gapfilling of Genome-Scale Metabolic Models"

#usr/bin/perl

use strict;
use LWP::UserAgent;

#written by: Wheaton Schroeder
#written to take a list of metabolite names, output list of kegg IDs

#read the file of names
open(METNAMES, "<met_names.txt") or die "unable to read file of metabolite names, reason: $!\n";
chomp(my @met_names = <METNAMES>);

#create file to output KEGGID list
open(MYKEDDIDS, ">kegg_ids.txt") or die "unable to create file to write kegg IDs to, reason: $!\n";

for(my $a = 0; $a <= $#met_names; $a++) {

	printf "working on metabolite \"%s\"\n", $met_names[$a];

	my $compartment = "[c]";

	#note if extracellular
	if($met_names[$a] =~ /\(Extracellular\)/) {
	
		#remove extracellular string, record new compartment
		$met_names[$a] =~ s/\(Extracellular\)//;
		$compartment = "[e]";
	
	}

	#remove E. coli note if present
	$met_names[$a] =~ s/\(E\.coli\) \*\*//g;

	#remove excess spaces
	$met_names[$a] =~ s/^\s+//;
	$met_names[$a] =~ s/\s+$//;
	
	printf "cleaned name: \"%s\"\n",$met_names[$a];

	#go to the link URL page to get the list of reactions for that map ID
	my $findCURL = "http://rest.kegg.jp/find/compound/".$met_names[$a]."/";
	#create a new user agent which pulls the KEGG API page
	my $UserAgent = LWP::UserAgent->new; #creates new user agent
	my $findCPage = $UserAgent->get($findCURL);
	$findCPage = $findCPage->content; 
	
	#split page into an array
	my @res_lines = split /\n/, $findCPage;
	
	#if there is only one result, accept it
	if($#res_lines eq 0) {
	
		printf "only one result\n";
	
		#now that we have the page number get each compound ID that is returned
		if($findCPage =~ /cpd:(C\d\d\d\d\d)/g) {
		
			printf MYKEDDIDS "%s%s",$1,$compartment;
			printf "found only kegg ID: %s%s\n", $1,$compartment;
		
		}
	
	} else {
	
		printf "multiple results\n";
		
		my $exact_match = 0;
		my $exact_id = "XX";
		my @matches = ( );
		
		#otherwise, if there are multiple results, search for exact text match
		for(my $b = 0; $b <= $#res_lines; $b++) {
	
			#split the multiple names
			my @pot_names = split /;/, $res_lines[$b];
			
			#check each name
			for(my $d = 0; $d <= $#pot_names; $d++) {
				
				#remove compound ID if first potential name
				$pot_names[$d] =~ s/^cpd\:C\d\d\d\d\d\s+//;
				
				#remove excess spaces
				$pot_names[$d] =~ s/^\s+//;
				$pot_names[$d] =~ s/\s+$//;
			
				#if exact match, say so and save the associated ID
				if($pot_names[$d] =~ /^$met_names[$a]$/i) {
				
					$exact_match = 1;
					
					if($res_lines[$b] =~ /cpd:(C\d\d\d\d\d)/g) {
					
						$exact_id = $1.$compartment;
					
					}
				
				}
			
			}
			
			if($res_lines[$b] =~ /cpd:(C\d\d\d\d\d)/g) {
				
				push @matches, $1.$compartment;
				
			}
			
		}
		
		if($exact_match eq 1) {
		
			#if an exact match was found report that ID only
			printf MYKEDDIDS "%s",$exact_id;
			printf "found exact kegg ID: %s\n",$exact_id;
		
		} else {
		
			#otherwise if no exact match, just list all potential matches
			for(my $c = 0; $c <= $#matches; $c++) {
			
				printf MYKEDDIDS "%s,",$matches[$c];
				printf "found potential kegg ID: %s\n",$matches[$c];
			
			}
			
		}
		
	
	}
	
	printf MYKEDDIDS "\n";
	printf "\n\n";

}
