## Supplemental File 3 for "A Novel Optimization-Based Tool to Automate Infeasible Cycle-Free Gapfilling of Genome-Scale Metabolic Models": FVA_combined_iJR904.gms

```
*GAMS code for Flux Variability Analysis of Arabidopsis Thaliana seed model
*Original code by: Mohammad Mazharul Islam
*modified for k-ath by: Wheaton Schroeder

$INLINECOM /*  */
$onlisting
$offdigit
$onempty

options limrow = 1000
	optCR = 1E-9
	optCA = 1E-9
	iterlim = 1000000
	decimals = 8
	reslim = 1000000
	work = 50000000
    lp = cplex;
        

SETS

	i						set of all metabolites
$include "all_mets_iJR904_iAF1260.txt"

	i_db(i)					set of all metabolites in the database 
$include "mets_iAF1260.txt"

	i_m(i)					set of all metabolites in the model
$include "mets_iJR904.txt"

	j						set of all reactions
$include "all_rxns_iJR904_iAF1260.txt"

	j_db(j)					set of all reactions in the database
$include "rxns_iAF1260.txt"

	j_m(j)					set of all reactions in the model
$include "rxns_iJR904.txt"

	exchange_m(j)				set of exchange reactions
$include "ex_rxns_iJR904.txt"

	exchange_db(j)				set of exchange reactions
$include "ex_rxns_iAF1260.txt"

	reg_off_m(j_m)	Reactions turned off (regulated)
$include "reg_rxns_iJR904.txt"

	reg_off_db(j_db)	Reactions turned off (regulated)
$include "reg_rxns_iAF1260.txt"


;

PARAMETERS

	UB(j) UB on rxns of kath 
  
	LB(j) LB on rxns of kath

	rxn_type_db(j_db)	reaction type of database reactions
$include "rxntype_iAF1260.txt"

	rxn_type_m(j_m)		reaction type of model reactions
$include "rxntype_iJR904.txt"

;

******************  Set the values of LB and UB ******************
SCALAR Vmax /10000/;

*** Set the experimental conditions

*irreversible reactions forward (rxntype = 1)
UB(j_db)$(rxn_type_db(j_db) = 1) = Vmax;
LB(j_db)$(rxn_type_db(j_db) = 1) = 0;

*For reversible reactions (rxntype = 0) 
UB(j_db)$(rxn_type_db(j_db) = 0) = Vmax;
LB(j_db)$(rxn_type_db(j_db) = 0) = -Vmax;

*irreversible reactions backwards (rxntype = -1)
UB(j_db)$(rxn_type_db(j_db) = -1) = 0;
LB(j_db)$(rxn_type_db(j_db) = -1) = -Vmax;

UB(j_db)$(reg_off_db(j_db)) = 0;
LB(j_db)$(reg_off_db(j_db)) = 0;

*irreversible reactions forward (rxntype = 1)
UB(j_m)$(rxn_type_m(j_m) = 1) = Vmax;
LB(j_m)$(rxn_type_m(j_m) = 1) = 0;

*For reversible reactions (rxntype = 0) 
UB(j_m)$(rxn_type_m(j_m) = 0) = Vmax;
LB(j_m)$(rxn_type_m(j_m) = 0) = -Vmax;

*irreversible reactions backwards (rxntype = -1)
UB(j_m)$(rxn_type_m(j_m) = -1) = 0;
LB(j_m)$(rxn_type_m(j_m) = -1) = -Vmax;

UB(j_m)$(reg_off_m(j_m)) = 0;
LB(j_m)$(reg_off_m(j_m)) = 0;

***** Set the conditions for the growth medium *****
*turn the growth medium back on only for iJR904 leave iAF1260 off
*first consider constant bounds
UB('EX_12ppd-S[e]') = Vmax;
UB('EX_15dap[e]') = Vmax;
UB('EX_26dap-M[e]') = Vmax;
UB('EX_2ddglcn[e]') = Vmax;
UB('EX_3hcinnm[e]') = Vmax;
UB('EX_3hpppn[e]') = Vmax;
UB('EX_4abut[e]') = Vmax;
UB('EX_acac[e]') = Vmax;
UB('EX_acald[e]') = Vmax;
UB('EX_acgam[e]') = Vmax;
UB('EX_acmana[e]') = Vmax;
UB('EX_acnam[e]') = Vmax;
UB('EX_ade[e]') = Vmax;
UB('EX_adn[e]') = Vmax;
UB('EX_ala-D[e]') = Vmax;
UB('EX_ala-L[e]') = Vmax;
UB('EX_alltn[e]') = Vmax;
UB('EX_amp[e]') = Vmax;
UB('EX_arab-L[e]') = Vmax;
UB('EX_arg-L[e]') = Vmax;
UB('EX_asn-L[e]') = Vmax;
UB('EX_asp-L[e]') = Vmax;
UB('EX_but[e]') = Vmax;
UB('EX_cbl1[e]') = Vmax;
UB('EX_chol[e]') = Vmax;
UB('EX_cit[e]') = Vmax;
UB('EX_crn[e]') = Vmax;
UB('EX_csn[e]') = Vmax;
UB('EX_cynt[e]') = Vmax;
UB('EX_cys-L[e]') = Vmax;
UB('EX_cytd[e]') = Vmax;
UB('EX_dad-2[e]') = Vmax;
UB('EX_dcyt[e]') = Vmax;
UB('EX_dgsn[e]') = Vmax;
UB('EX_dha[e]') = Vmax;
UB('EX_din[e]') = Vmax;
UB('EX_dms[e]') = Vmax;
UB('EX_dmso[e]') = Vmax;
UB('EX_duri[e]') = Vmax;
UB('EX_etoh[e]') = Vmax;
UB('EX_for[e]') = Vmax;
UB('EX_fru[e]') = Vmax;
UB('EX_fuc-L[e]') = Vmax;
UB('EX_fuc1p-L[e]') = Vmax;
UB('EX_fum[e]') = Vmax;
UB('EX_g6p[e]') = Vmax;
UB('EX_gal[e]') = Vmax;
UB('EX_galct-D[e]') = Vmax;
UB('EX_galctn-D[e]') = Vmax;
UB('EX_galt[e]') = Vmax;
UB('EX_galur[e]') = Vmax;
UB('EX_gam[e]') = Vmax;
UB('EX_gbbtn[e]') = Vmax;
UB('EX_glcn[e]') = Vmax;
UB('EX_glcr[e]') = Vmax;
UB('EX_glcur[e]') = Vmax;
UB('EX_gln-L[e]') = Vmax;
UB('EX_glu-L[e]') = Vmax;
UB('EX_gly[e]') = Vmax;
UB('EX_glyald[e]') = Vmax;
UB('EX_glyb[e]') = Vmax;
UB('EX_glyc3p[e]') = Vmax;
UB('EX_glyclt[e]') = Vmax;
UB('EX_gsn[e]') = Vmax;
UB('EX_gua[e]') = Vmax;
UB('EX_hdca[e]') = Vmax;
UB('EX_his-L[e]') = Vmax;
UB('EX_hxan[e]') = Vmax;
UB('EX_idon-L[e]') = Vmax;
UB('EX_ile-L[e]') = Vmax;
UB('EX_indole[e]') = Vmax;
UB('EX_ins[e]') = Vmax;
UB('EX_lcts[e]') = Vmax;
UB('EX_leu-L[e]') = Vmax;
UB('EX_lys-L[e]') = Vmax;
UB('EX_malt[e]') = Vmax;
UB('EX_malthx[e]') = Vmax;
UB('EX_maltpt[e]') = Vmax;
UB('EX_malttr[e]') = Vmax;
UB('EX_maltttr[e]') = Vmax;
UB('EX_man[e]') = Vmax;
UB('EX_man6p[e]') = Vmax;
UB('EX_melib[e]') = Vmax;
UB('EX_met-D[e]') = Vmax;
UB('EX_met-L[e]') = Vmax;
UB('EX_mnl[e]') = Vmax;
UB('EX_nac[e]') = Vmax;
UB('EX_nad[e]') = Vmax;
UB('EX_nmn[e]') = Vmax;
UB('EX_no2[e]') = Vmax;
UB('EX_no3[e]') = Vmax;
UB('EX_ocdca[e]') = Vmax;
UB('EX_orn[e]') = Vmax;
UB('EX_phe-L[e]') = Vmax;
UB('EX_pnto-R[e]') = Vmax;
UB('EX_pppn[e]') = Vmax;
UB('EX_pro-L[e]') = Vmax;
UB('EX_ptrc[e]') = Vmax;
UB('EX_rib-D[e]') = Vmax;
UB('EX_rmn[e]') = Vmax;
UB('EX_sbt-D[e]') = Vmax;
UB('EX_ser-D[e]') = Vmax;
UB('EX_ser-L[e]') = Vmax;
UB('EX_spmd[e]') = Vmax;
UB('EX_sucr[e]') = Vmax;
UB('EX_tartr-L[e]') = Vmax;
UB('EX_taur[e]') = Vmax;
UB('EX_thm[e]') = Vmax;
UB('EX_thr-L[e]') = Vmax;
UB('EX_thymd[e]') = Vmax;
UB('EX_tma[e]') = Vmax;
UB('EX_tmao[e]') = Vmax;
UB('EX_tre[e]') = Vmax;
UB('EX_trp-L[e]') = Vmax;
UB('EX_tsul[e]') = Vmax;
UB('EX_ttdca[e]') = Vmax;
UB('EX_tyr-L[e]') = Vmax;
UB('EX_ura[e]') = Vmax;
UB('EX_urea[e]') = Vmax;
UB('EX_uri[e]') = Vmax;
UB('EX_val-L[e]') = Vmax;
UB('EX_xan[e]') = Vmax;
UB('EX_xtsn[e]') = Vmax;
UB('EX_xyl-D[e]') = Vmax;
UB('EX_co2[e]') = Vmax;
UB('EX_fe2[e]') = Vmax;
UB('EX_h[e]') = Vmax;
UB('EX_h2o[e]') = Vmax;
UB('EX_k[e]') = Vmax;
UB('EX_na1[e]') = Vmax;
UB('EX_nh4[e]') = Vmax;
UB('EX_pi[e]') = Vmax;
UB('EX_so4[e]') = Vmax;

LB('EX_12ppd-S[e]') = 0;
LB('EX_15dap[e]') = 0;
LB('EX_26dap-M[e]') = 0;
LB('EX_2ddglcn[e]') = 0;
LB('EX_3hcinnm[e]') = 0;
LB('EX_3hpppn[e]') = 0;
LB('EX_4abut[e]') = 0;
LB('EX_acac[e]') = 0;
LB('EX_acald[e]') = 0;
LB('EX_acgam[e]') = 0;
LB('EX_acmana[e]') = 0;
LB('EX_acnam[e]') = 0;
LB('EX_ade[e]') = 0;
LB('EX_adn[e]') = 0;
LB('EX_ala-D[e]') = 0;
LB('EX_ala-L[e]') = 0;
LB('EX_alltn[e]') = 0;
LB('EX_amp[e]') = 0;
LB('EX_arab-L[e]') = 0;
LB('EX_arg-L[e]') = 0;
LB('EX_asn-L[e]') = 0;
LB('EX_asp-L[e]') = 0;
LB('EX_but[e]') = 0;
LB('EX_cbl1[e]') = 0;
LB('EX_chol[e]') = 0;
LB('EX_cit[e]') = 0;
LB('EX_crn[e]') = 0;
LB('EX_csn[e]') = 0;
LB('EX_cynt[e]') = 0;
LB('EX_cys-L[e]') = 0;
LB('EX_cytd[e]') = 0;
LB('EX_dad-2[e]') = 0;
LB('EX_dcyt[e]') = 0;
LB('EX_dgsn[e]') = 0;
LB('EX_dha[e]') = 0;
LB('EX_din[e]') = 0;
LB('EX_dms[e]') = 0;
LB('EX_dmso[e]') = 0;
LB('EX_duri[e]') = 0;
LB('EX_etoh[e]') = 0;
LB('EX_for[e]') = 0;
LB('EX_fru[e]') = 0;
LB('EX_fuc-L[e]') = 0;
LB('EX_fuc1p-L[e]') = 0;
LB('EX_fum[e]') = 0;
LB('EX_g6p[e]') = 0;
LB('EX_gal[e]') = 0;
LB('EX_galct-D[e]') = 0;
LB('EX_galctn-D[e]') = 0;
LB('EX_galt[e]') = 0;
LB('EX_galur[e]') = 0;
LB('EX_gam[e]') = 0;
LB('EX_gbbtn[e]') = 0;
LB('EX_glcn[e]') = 0;
LB('EX_glcr[e]') = 0;
LB('EX_glcur[e]') = 0;
LB('EX_gln-L[e]') = 0;
LB('EX_glu-L[e]') = 0;
LB('EX_gly[e]') = 0;
LB('EX_glyald[e]') = 0;
LB('EX_glyb[e]') = 0;
LB('EX_glyc3p[e]') = 0;
LB('EX_glyclt[e]') = 0;
LB('EX_gsn[e]') = 0;
LB('EX_gua[e]') = 0;
LB('EX_hdca[e]') = 0;
LB('EX_his-L[e]') = 0;
LB('EX_hxan[e]') = 0;
LB('EX_idon-L[e]') = 0;
LB('EX_ile-L[e]') = 0;
LB('EX_indole[e]') = 0;
LB('EX_ins[e]') = 0;
LB('EX_lcts[e]') = 0;
LB('EX_leu-L[e]') = 0;
LB('EX_lys-L[e]') = 0;
LB('EX_malt[e]') = 0;
LB('EX_malthx[e]') = 0;
LB('EX_maltpt[e]') = 0;
LB('EX_malttr[e]') = 0;
LB('EX_maltttr[e]') = 0;
LB('EX_man[e]') = 0;
LB('EX_man6p[e]') = 0;
LB('EX_melib[e]') = 0;
LB('EX_met-D[e]') = 0;
LB('EX_met-L[e]') = 0;
LB('EX_mnl[e]') = 0;
LB('EX_nac[e]') = 0;
LB('EX_nad[e]') = 0;
LB('EX_nmn[e]') = 0;
LB('EX_no2[e]') = 0;
LB('EX_no3[e]') = 0;
LB('EX_ocdca[e]') = 0;
LB('EX_orn[e]') = 0;
LB('EX_phe-L[e]') = 0;
LB('EX_pnto-R[e]') = 0;
LB('EX_pppn[e]') = 0;
LB('EX_pro-L[e]') = 0;
LB('EX_ptrc[e]') = 0;
LB('EX_rib-D[e]') = 0;
LB('EX_rmn[e]') = 0;
LB('EX_sbt-D[e]') = 0;
LB('EX_ser-D[e]') = 0;
LB('EX_ser-L[e]') = 0;
LB('EX_spmd[e]') = 0;
LB('EX_sucr[e]') = 0;
LB('EX_tartr-L[e]') = 0;
LB('EX_taur[e]') = 0;
LB('EX_thm[e]') = 0;
LB('EX_thr-L[e]') = 0;
LB('EX_thymd[e]') = 0;
LB('EX_tma[e]') = 0;
LB('EX_tmao[e]') = 0;
LB('EX_tre[e]') = 0;
LB('EX_trp-L[e]') = 0;
LB('EX_tsul[e]') = 0;
LB('EX_ttdca[e]') = 0;
LB('EX_tyr-L[e]') = 0;
LB('EX_ura[e]') = 0;
LB('EX_urea[e]') = 0;
LB('EX_uri[e]') = 0;
LB('EX_val-L[e]') = 0;
LB('EX_xan[e]') = 0;
LB('EX_xtsn[e]') = 0;
LB('EX_xyl-D[e]') = 0;
LB('EX_co2[e]') = -Vmax;
LB('EX_fe2[e]') = -Vmax;
LB('EX_h[e]') = -Vmax;
LB('EX_h2o[e]') = -Vmax;
LB('EX_k[e]') = -Vmax;
LB('EX_na1[e]') = -Vmax;
LB('EX_nh4[e]') = -Vmax;
LB('EX_pi[e]') = -Vmax;
LB('EX_so4[e]') = -Vmax;

LB(exchange_db) = 0;
UB(exchange_db) = 0;

*now consider variable bounds
*aerobic conditions
UB('EX_ac[e]') = Vmax;
UB('EX_akg[e]') = Vmax;
UB('EX_glc[e]') = Vmax;
UB('EX_glyc[e]') = Vmax;
UB('EX_lac-D[e]') = Vmax;
UB('EX_lac-L[e]') = Vmax;
UB('EX_mal-L[e]') = Vmax;
UB('EX_o2[e]') = 0;
UB('EX_pyr[e]') = Vmax;
UB('EX_succ[e]') = Vmax;

LB('EX_ac[e]') = -5;
LB('EX_akg[e]') = 0;
LB('EX_glc[e]') = 0;
LB('EX_glyc[e]') = 0;
LB('EX_lac-D[e]') = 0;
LB('EX_lac-L[e]') = 0;
LB('EX_mal-L[e]') = 0;
LB('EX_o2[e]') = -20;
LB('EX_pyr[e]') = 0;
LB('EX_succ[e]') = 0;

****************stoichiometry*****************
PARAMETERS
  Skath(i,j) Stoichiometric matrix for kath 
$include "Sij_iJR904_iAF1260.txt"
;

PARAMETERS
C(j)
max(j)
min(j);
alias (j1kath, j);

VARIABLES
	zprimal_kath     primal objective function (rxn fluxes in each loop) for kath
    vkath(j)       Flux of kath rxns
;
	

vkath.lo(j)=LB(j);
vkath.up(j)=UB(j);


*********************EQUATIONS NAMES**************************
EQUATIONS

	primalobj_kath			primal objective function (rxn fluxes in each loop) for kath
	massbalance_kath(i)  	Mass balance for kath
	
;

****************************** Equations*******************************
primalobj_kath..		  zprimal_kath =e= sum(j$(C(j) eq 1),vkath(j));

massbalance_kath(i)..     sum(j,Skath(i,j)*vkath(j)) =e= 0;
*****************************************************************************

file result /FVA_result_iJR904_combined.txt/;
put result;
result.lw = 24;

put 'FVA results for kath'/;
put 'Reaction','                        MIN','         MAX','         LB','          UB','       Model Status'/;
put '---------------------------------------------------------------------------------------------------'/;

model primalkath
/
primalobj_kath
massbalance_kath
/
;

LOOP(j1kath,

	C(j) = 0;
	C(j1kath) = 1;
	
	Solve primalkath using lp maximizing zprimal_kath;
	max(j1kath) = zprimal_kath.l;

	Solve primalkath using lp minimizing zprimal_kath;
	min(j1kath) = zprimal_kath.l;
	put j1kath.tl,min(j1kath),max(j1kath),LB(j1kath),UB(j1kath),primalkath.modelstat/;

);

put //;

put 'Fluxes hitting upper bounds'/;
put '------------------------------------------------------------------------'/;

LOOP(j,
if(max(j)>= 9000, put j.tl,'	Max flux =', max(j)/;
);
);

put 'Fluxes hitting lower bounds'/;
put '------------------------------------------------------------------------'/;
LOOP(j,
if(min(j)<= -9000, put j.tl,'	Min flux =', min(j)/;
);
);

put //;

put 'Blocked reactions'/;
put '------------------------------------------------------------------------'/;
LOOP(j,
if(bool_and(max(j)= 0,min(j)= 0), put j.tl/;
);
);
```
