## Supplemental File 3 for "A Novel Optimization-Based Tool to Automate Infeasible Cycle-Free Gapfilling of Genome-Scale Metabolic Models": get_solve_times.pl

```
#usr/bin/perl

use strict;

#read the output file of OptFill as the input, assign to array
open(INPUT, "<optfill_iJR904_iAF1260.txt") or die "could not open input file, reason: $!\n";
chomp(my @input = <INPUT>);

#have output file to write solution times
open(OUTPUT1, ">solutions_CPs.csv") or die "could not create/write solution times file, reason: $!\n";

#have an output for TFP statistics
open(OUTPUT2, ">solutions_TICs.csv") or die "could not create/write TIC solution information file, reason: $!\n";

for(my $a = 0; $a <= $#input; $a++) {
	
	if($input[$a] =~ /TIC NUMBER\s+(\d+)\.\d+/) {
	
		printf OUTPUT2 "%s,", $1;
	
	}
	
	if($input[$a] =~ /Number of Reactions\:\s+(\d+)\.\d+/) {
	
		printf OUTPUT2 "%s,", $1;
	
	}
	
	if($input[$a] =~ /solution\stime\:\s+(\d+\.\d+)/) {
	
		printf OUTPUT2 "%s\n", $1;
	
	}
	
	if($input[$a] =~ /OUTER OBJECTIVE VALUE\:\s+(\d+)\.\d+/) {
	
		printf OUTPUT1 "%s,", $1;
	
	}
	
	if($input[$a] =~ /INNER OBJECTIVE VALUE 1\:\s+(\d+)\.\d+/) {
	
		printf OUTPUT1 "%s,", $1;
	
	}
	
	if($input[$a] =~ /INNER OBJECTIVE VALUE 2\:\s+(\d+)\.\d+/) {
	
		printf OUTPUT1 "%s,", $1;
	
	}
	
	if($input[$a] =~ /new\smodel\sbiomass\srate:\s+(\d+\.\d+)/) {
	
		printf OUTPUT1 "%s,", $1;
	
	}
	
	if($input[$a] =~ /solve\stime\:\s+(\d+\.\d+)/) {
	
		printf OUTPUT1 "%s\n", $1;
	
	}	
	
}
```
