## Supplemental File 3 for "A Novel Optimization-Based Tool to Automate Infeasible Cycle-Free Gapfilling of Genome-Scale Metabolic Models": make_m_and_db.pl

```
#usr/bin/perl

#Written by: Wheaton Schroeder
#Latest version: 9/4/19
use strict;

#oepn the list of all reactions
open(MANDDB, "<test_3_new_rxns.txt") or die "could not open list of new reactions file, reason: $!\n";
chomp(my @new_rxns = <MANDDB>);

#creat files to write database and model files
open(ADDTODB, ">test_3_add_to_db.txt") or die "could not write add to database file, reason: $!\n";
open(ADDTOM, ">test_3_add_to_m.txt") or die "could not write add to model file, reason: $!\n";

#randomly assign value of 0 to 1 to each reaction, sort to database or model based on value
for(my $a = 0; $a <= $#new_rxns; $a++) {

	my $temp_rand = rand();
	
	if ($temp_rand le 0.8) {
	
		#will add to new database
		printf ADDTOM "%s\n", $new_rxns[$a];
	
	} else {
	
		#will add to new model
		printf ADDTODB "%s\n", $new_rxns[$a];
	
	}

}
```
