## Supplemental File 3 for "A Novel Optimization-Based Tool to Automate Infeasible Cycle-Free Gapfilling of Genome-Scale Metabolic Models": OptFill_iJR904_iAF1260.gms

```
****************************************************************************************
*                                    OptFill                                           *
*                Created and written by: Wheaton Schroeder                             *
*Created to fill in a model from a database using reversible and irreversible adds. As *
*this formulation has significantly more variables (especially binary variables) than a*
*Gapfill approach the idea of this problem/formulation/code/script is to use the       *
*results of gapfill (where individual solution sets do not cause an issue like a TIC,  *
*these sets being on a per-metabolite basis) and tries to use the gapfill solutions to *
*fix as many metabolites as possible without causing a TIC, sometimes by fixing direc- *
*tionality when possible.                                                              *
*Latest version: 9/23/19                                                              *
*Generally named "optfill.gms"														   *
****************************************************************************************

$INLINECOM /*  */
$offlisting
$onempty

OPTIONS

	decimals = 8
	solprint = on
	mip = cplex
	minlp = baron
	
;

*define sets
SETS

	i						set of all metabolites
$include "all_mets_iJR904_iAF1260.txt"

	i_db(i)					set of all metabolites in the database 
$include "mets_iAF1260.txt"

	i_m(i)					set of all metabolites in the model
$include "mets_iJR904.txt"

	biomass_precursors(i)	set of metabolites which are the precursors to biomass
$include "biomass_precursors_iJR904.txt"

	j						set of all reactions
$include "all_rxns_iJR904_iAF1260.txt"

	j_db(j)					set of all reactions in the database
$include "rxns_iAF1260.txt"

	j_m(j)					set of all reactions in the model
$include "rxns_iJR904.txt"

	p /1*10000/				set of TICs
	
	eqncounter(p)			counts the number of TIC found to present
	
	eqncounter_2(p)			counter number of unique connecting solutions present
	
	eqncounter_3(p)			counter number of unique reversible connecting solutions present

	exchange(j)				set of exchange reactions
$include "ex_rxns_iJR904.txt"

	reg_off_m(j_m)	Reactions turned off (regulated)
$include "reg_rxns_iJR904.txt"

	reg_off_db(j_db)	Reactions turned off (regulated)
$include "reg_rxns_iAF1260.txt"

;

*initially, there are not previous solutions
eqncounter(p)=no;
eqncounter_2(p)=no;
eqncounter_3(p)=no;

*define parameters
PARAMETERS

	S(i,j)				Sij matrix for the database and model
$include "Sij_iJR904_iAF1260.txt"

	rxn_type_db(j_db)	reaction type of database reactions
$include "rxntype_iAF1260.txt"

	rxn_type_m(j_m)		reaction type of model reactions
$include "rxntype_iJR904.txt"

	vUB(j)				set of upper bound of flux for reaction j
	
	vLB(j)				set of lower bound of flux for reaction j

	z_max				maximum value of z
	
	temp				temporary value for increasing other parameters incrementally 
	
	temp1				temporary value for increasing other parameters incrementally 
	
	temp2				temporary value for increasing other parameters incrementally 
	
	TIC_count			keep a count of the number of TICs found
	
	TIC_count_temp		this is an updater parameter for the number of TICs found

	TFP_soln(j,p)		stores non-directional previous solution - eta
	
	TFP_soln_f(j,p)		stores reactions taking place in the forward direction in previous normal solutions to the TIC finding problem
	
	TFP_soln_b(j,p)		stores reactions taking place in the forward direction in previous normal solutions to the TIC finding problem
	
	done				stores wheter or not all possible TICs have been found
	
	count_rxns			counts the number of reactions proposing to be added
	
	count_rxns_temp		counts the number of reactions proposing to be added=
	
	phi					maximum number of rxns in the current TIC
	
	phi_temp			updater variable of phi
	
	phi_max				maximum size of any TIC
	
	CPs_soln(j_db,p)	stores the set of reactions used to gapfill the model in a given solution iteration undirectional
	
	CPs_soln_f(j_db,p)	stores the set of reactions used to gapfill the model in a given solution iteration stores forward
	
	CPs_soln_b(j_db,p)	stores the set of reactions used to gapfill the model in a given solution iteration stores backward
	
	num_omegas(p)		stores the number of omegas assoiated with a particular solution helps avoid repeat solutions
	
	curr_CPs_soln_f		stores the delta value for the current connecting solution
	
	curr_CPs_soln_b		stores the rho value for the current connecting solution
	
	Num_Solns			number of connecting solutions identified
	
	bio_rate(p)			stores the biomass growth rate associated with connecting solution p will be used to perform statistics on it
	
	bio_mean			reports mean biomass growth rate of connecting solutions
	
	bio_min				reports minimum biomass growth rate of connecting solutions
	
	bio_max				reports maximum biomass growth rate of connecting solutions
	
	bio_sd				reports standard deviation growth rate of connecting solutions
	
	iter_max			maximum number of iterations for error handling
	
	iter				count of iterations used for error handling
	
	iter_CP1			stores number of interations for error handling used by CP1
	
	iter_CP2			stores number of interations for error handling used by CP2
	
	iter_CP3			stores number of interations for error handling used by CP3
	
	TFP_rate(p)			stores the solution time of the TIC finding problems
	
	TFP_mean			reports mean TIC finding problem solution time
	
	TFP_min				reports minimum TIC finding problem solution time
	
	TFP_max				reports maximum TIC finding problem solution time
	
	TFP_sd				reports standard deviation TIC finding problem solution time
	
	time_cp1			time to solve CP1
	
	time_cp2			time to solve CP2
	
	time_cp3			time to solve CP3
	
	CP_rate(p)			stores the solution time of the connecting problems
	
	CP_mean				reports mean connecting problems solution time
	
	CP_min				reports minimum connecting problems solution time
	
	CP_max				reports maximum connecting problems solution time
	
	CP_sd				reports standard deviation connecting problems solution time
	
	base_prod(i_m)		stores if base model can produce a given metabolite
	
	xi(i,j)				stores 1 if metabolite i is on the rhs of reaction j zero otherwise
	
	psi(i,j) 			stores 1 if metabolite i is on the lhs of reaction j zero otherwise
	
;

xi(i,j) = 0;
xi(i,j)$(S(i,j) > 0) = 1;
psi(i,j) = 0;
psi(i,j)$(S(i,j) < 0) = 1;

*no solutions initially
TFP_soln(j,p)=no;
TFP_soln_f(j,p)=no;
TFP_soln_b(j,p)=no;
CPs_soln(j_db,p)=no;
CPs_soln_f(j_db,p)=no;
CPs_soln_b(j_db,p)=no;
num_omegas(p)=no;

Num_Solns = 0;

*loop is not done yet
done = 0;

*set values of big M and epsilon
SCALAR M /1E3/;
SCALAR epsilon /1E-3/;

*default set bounds to +-M
vLB(j) = -M;
vUB(j) = M;

*set reaction bounds
* irreversible reactions, forwards
vLB(j_m)$(rxn_type_m(j_m) eq 1) = 0;
vUB(j_m)$(rxn_type_m(j_m) eq 1) = M;

* Reversible reactions
vLB(j_m)$(rxn_type_m(j_m) eq 0) = -M;
vUB(j_m)$(rxn_type_m(j_m) eq 0) = M;

* irreversible reactions, backwards
vLB(j_m)$(rxn_type_m(j_m) eq -1) = -M;
vUB(j_m)$(rxn_type_m(j_m) eq -1) = 0;

* reaction bounds for database
vLB(j_db)$(rxn_type_db(j_db) eq 1) = 0;
vUB(j_db)$(rxn_type_db(j_db) eq 1) = M;

vLB(j_db)$(rxn_type_db(j_db) eq 0) = -M;
vUB(j_db)$(rxn_type_db(j_db) eq 0) = M;

vLB(j_db)$(rxn_type_db(j_db) eq -1) = -M;
vUB(j_db)$(rxn_type_db(j_db) eq -1) = 0;

vLB(reg_off_m) = 0;
vLB(reg_off_db) = 0;
vUB(reg_off_m) = 0;
vUB(reg_off_db) = 0;

**************************** TIC finding Problem ************************************

*define the variables for the TIC finding problem
BINARY VARIABLES

	eta(j)				has a value of 1 if reaction j participates in current TIC zero otherwise
	alpha(j)			if v(j) is positive and eta equals 1 then this has a value of 1 indicates positive participation in the TIC
	beta(j)				if v(j) is negative and eta equals 1 then this has a value of 1 indicates negative participation in the TIC
	gamma(p)			part of the integer cuts allows for reverse direction TICs to be identified
	
VARIABLES

	E					objective variable
	v(j)				flux values associated with the TIC
	
;

*define the constraint equations
EQUATIONS

	TFP_obj
	TFP_const_1(j)
	TFP_const_2(j)
	TFP_const_3(j)
	TFP_const_4(j)
	TFP_const_5(i)
	TFP_const_6
	TFP_const_7
	TFP_const_8(j)
	TFP_const_9(j)
	TFP_const_10(j)
	TIC_int_cut_1(p)
	TIC_int_cut_2(p)

;

phi = 1;
*phi_max = sum(j, 1);
phi_max = 9;

*turn off growth medium when checking for TICs
vLB(exchange) = 0;
vUB(exchange) = 0;

*turn the growth medium back on only for iJR904 leave iAF1260 off
*first consider constant bounds
vUB('EX_12ppd-S[e]') = 0;
vUB('EX_15dap[e]') = 0;
vUB('EX_26dap-M[e]') = 0;
vUB('EX_2ddglcn[e]') = 0;
vUB('EX_3hcinnm[e]') = 0;
vUB('EX_3hpppn[e]') = 0;
vUB('EX_4abut[e]') = 0;
vUB('EX_acac[e]') = 0;
vUB('EX_acald[e]') = 0;
vUB('EX_acgam[e]') = 0;
vUB('EX_acmana[e]') = 0;
vUB('EX_acnam[e]') = 0;
vUB('EX_ade[e]') = 0;
vUB('EX_adn[e]') = 0;
vUB('EX_ala-D[e]') = 0;
vUB('EX_ala-L[e]') = 0;
vUB('EX_alltn[e]') = 0;
vUB('EX_amp[e]') = 0;
vUB('EX_arab-L[e]') = 0;
vUB('EX_arg-L[e]') = 0;
vUB('EX_asn-L[e]') = 0;
vUB('EX_asp-L[e]') = 0;
vUB('EX_but[e]') = 0;
vUB('EX_cbl1[e]') = 0;
vUB('EX_chol[e]') = 0;
vUB('EX_cit[e]') = 0;
vUB('EX_crn[e]') = 0;
vUB('EX_csn[e]') = 0;
vUB('EX_cynt[e]') = 0;
vUB('EX_cys-L[e]') = 0;
vUB('EX_cytd[e]') = 0;
vUB('EX_dad-2[e]') = 0;
vUB('EX_dcyt[e]') = 0;
vUB('EX_dgsn[e]') = 0;
vUB('EX_dha[e]') = 0;
vUB('EX_din[e]') = 0;
vUB('EX_dms[e]') = 0;
vUB('EX_dmso[e]') = 0;
vUB('EX_duri[e]') = 0;
vUB('EX_etoh[e]') = 0;
vUB('EX_for[e]') = 0;
vUB('EX_fru[e]') = 0;
vUB('EX_fuc-L[e]') = 0;
vUB('EX_fuc1p-L[e]') = 0;
vUB('EX_fum[e]') = 0;
vUB('EX_g6p[e]') = 0;
vUB('EX_gal[e]') = 0;
vUB('EX_galct-D[e]') = 0;
vUB('EX_galctn-D[e]') = 0;
vUB('EX_galt[e]') = 0;
vUB('EX_galur[e]') = 0;
vUB('EX_gam[e]') = 0;
vUB('EX_gbbtn[e]') = 0;
vUB('EX_glcn[e]') = 0;
vUB('EX_glcr[e]') = 0;
vUB('EX_glcur[e]') = 0;
vUB('EX_gln-L[e]') = 0;
vUB('EX_glu-L[e]') = 0;
vUB('EX_gly[e]') = 0;
vUB('EX_glyald[e]') = 0;
vUB('EX_glyb[e]') = 0;
vUB('EX_glyc3p[e]') = 0;
vUB('EX_glyclt[e]') = 0;
vUB('EX_gsn[e]') = 0;
vUB('EX_gua[e]') = 0;
vUB('EX_hdca[e]') = 0;
vUB('EX_his-L[e]') = 0;
vUB('EX_hxan[e]') = 0;
vUB('EX_idon-L[e]') = 0;
vUB('EX_ile-L[e]') = 0;
vUB('EX_indole[e]') = 0;
vUB('EX_ins[e]') = 0;
vUB('EX_lcts[e]') = 0;
vUB('EX_leu-L[e]') = 0;
vUB('EX_lys-L[e]') = 0;
vUB('EX_malt[e]') = 0;
vUB('EX_malthx[e]') = 0;
vUB('EX_maltpt[e]') = 0;
vUB('EX_malttr[e]') = 0;
vUB('EX_maltttr[e]') = 0;
vUB('EX_man[e]') = 0;
vUB('EX_man6p[e]') = 0;
vUB('EX_melib[e]') = 0;
vUB('EX_met-D[e]') = 0;
vUB('EX_met-L[e]') = 0;
vUB('EX_mnl[e]') = 0;
vUB('EX_nac[e]') = 0;
vUB('EX_nad[e]') = 0;
vUB('EX_nmn[e]') = 0;
vUB('EX_no2[e]') = 0;
vUB('EX_no3[e]') = 0;
vUB('EX_ocdca[e]') = 0;
vUB('EX_orn[e]') = 0;
vUB('EX_phe-L[e]') = 0;
vUB('EX_pnto-R[e]') = 0;
vUB('EX_pppn[e]') = 0;
vUB('EX_pro-L[e]') = 0;
vUB('EX_ptrc[e]') = 0;
vUB('EX_rib-D[e]') = 0;
vUB('EX_rmn[e]') = 0;
vUB('EX_sbt-D[e]') = 0;
vUB('EX_ser-D[e]') = 0;
vUB('EX_ser-L[e]') = 0;
vUB('EX_spmd[e]') = 0;
vUB('EX_sucr[e]') = 0;
vUB('EX_tartr-L[e]') = 0;
vUB('EX_taur[e]') = 0;
vUB('EX_thm[e]') = 0;
vUB('EX_thr-L[e]') = 0;
vUB('EX_thymd[e]') = 0;
vUB('EX_tma[e]') = 0;
vUB('EX_tmao[e]') = 0;
vUB('EX_tre[e]') = 0;
vUB('EX_trp-L[e]') = 0;
vUB('EX_tsul[e]') = 0;
vUB('EX_ttdca[e]') = 0;
vUB('EX_tyr-L[e]') = 0;
vUB('EX_ura[e]') = 0;
vUB('EX_urea[e]') = 0;
vUB('EX_uri[e]') = 0;
vUB('EX_val-L[e]') = 0;
vUB('EX_xan[e]') = 0;
vUB('EX_xtsn[e]') = 0;
vUB('EX_xyl-D[e]') = 0;
vUB('EX_co2[e]') = 0;
vUB('EX_fe2[e]') = 0;
vUB('EX_h[e]') = 0;
vUB('EX_h2o[e]') = 0;
vUB('EX_k[e]') = 0;
vUB('EX_na1[e]') = 0;
vUB('EX_nh4[e]') = 0;
vUB('EX_pi[e]') = 0;
vUB('EX_so4[e]') = 0;

vLB('EX_12ppd-S[e]') = 0;
vLB('EX_15dap[e]') = 0;
vLB('EX_26dap-M[e]') = 0;
vLB('EX_2ddglcn[e]') = 0;
vLB('EX_3hcinnm[e]') = 0;
vLB('EX_3hpppn[e]') = 0;
vLB('EX_4abut[e]') = 0;
vLB('EX_acac[e]') = 0;
vLB('EX_acald[e]') = 0;
vLB('EX_acgam[e]') = 0;
vLB('EX_acmana[e]') = 0;
vLB('EX_acnam[e]') = 0;
vLB('EX_ade[e]') = 0;
vLB('EX_adn[e]') = 0;
vLB('EX_ala-D[e]') = 0;
vLB('EX_ala-L[e]') = 0;
vLB('EX_alltn[e]') = 0;
vLB('EX_amp[e]') = 0;
vLB('EX_arab-L[e]') = 0;
vLB('EX_arg-L[e]') = 0;
vLB('EX_asn-L[e]') = 0;
vLB('EX_asp-L[e]') = 0;
vLB('EX_but[e]') = 0;
vLB('EX_cbl1[e]') = 0;
vLB('EX_chol[e]') = 0;
vLB('EX_cit[e]') = 0;
vLB('EX_crn[e]') = 0;
vLB('EX_csn[e]') = 0;
vLB('EX_cynt[e]') = 0;
vLB('EX_cys-L[e]') = 0;
vLB('EX_cytd[e]') = 0;
vLB('EX_dad-2[e]') = 0;
vLB('EX_dcyt[e]') = 0;
vLB('EX_dgsn[e]') = 0;
vLB('EX_dha[e]') = 0;
vLB('EX_din[e]') = 0;
vLB('EX_dms[e]') = 0;
vLB('EX_dmso[e]') = 0;
vLB('EX_duri[e]') = 0;
vLB('EX_etoh[e]') = 0;
vLB('EX_for[e]') = 0;
vLB('EX_fru[e]') = 0;
vLB('EX_fuc-L[e]') = 0;
vLB('EX_fuc1p-L[e]') = 0;
vLB('EX_fum[e]') = 0;
vLB('EX_g6p[e]') = 0;
vLB('EX_gal[e]') = 0;
vLB('EX_galct-D[e]') = 0;
vLB('EX_galctn-D[e]') = 0;
vLB('EX_galt[e]') = 0;
vLB('EX_galur[e]') = 0;
vLB('EX_gam[e]') = 0;
vLB('EX_gbbtn[e]') = 0;
vLB('EX_glcn[e]') = 0;
vLB('EX_glcr[e]') = 0;
vLB('EX_glcur[e]') = 0;
vLB('EX_gln-L[e]') = 0;
vLB('EX_glu-L[e]') = 0;
vLB('EX_gly[e]') = 0;
vLB('EX_glyald[e]') = 0;
vLB('EX_glyb[e]') = 0;
vLB('EX_glyc3p[e]') = 0;
vLB('EX_glyclt[e]') = 0;
vLB('EX_gsn[e]') = 0;
vLB('EX_gua[e]') = 0;
vLB('EX_hdca[e]') = 0;
vLB('EX_his-L[e]') = 0;
vLB('EX_hxan[e]') = 0;
vLB('EX_idon-L[e]') = 0;
vLB('EX_ile-L[e]') = 0;
vLB('EX_indole[e]') = 0;
vLB('EX_ins[e]') = 0;
vLB('EX_lcts[e]') = 0;
vLB('EX_leu-L[e]') = 0;
vLB('EX_lys-L[e]') = 0;
vLB('EX_malt[e]') = 0;
vLB('EX_malthx[e]') = 0;
vLB('EX_maltpt[e]') = 0;
vLB('EX_malttr[e]') = 0;
vLB('EX_maltttr[e]') = 0;
vLB('EX_man[e]') = 0;
vLB('EX_man6p[e]') = 0;
vLB('EX_melib[e]') = 0;
vLB('EX_met-D[e]') = 0;
vLB('EX_met-L[e]') = 0;
vLB('EX_mnl[e]') = 0;
vLB('EX_nac[e]') = 0;
vLB('EX_nad[e]') = 0;
vLB('EX_nmn[e]') = 0;
vLB('EX_no2[e]') = 0;
vLB('EX_no3[e]') = 0;
vLB('EX_ocdca[e]') = 0;
vLB('EX_orn[e]') = 0;
vLB('EX_phe-L[e]') = 0;
vLB('EX_pnto-R[e]') = 0;
vLB('EX_pppn[e]') = 0;
vLB('EX_pro-L[e]') = 0;
vLB('EX_ptrc[e]') = 0;
vLB('EX_rib-D[e]') = 0;
vLB('EX_rmn[e]') = 0;
vLB('EX_sbt-D[e]') = 0;
vLB('EX_ser-D[e]') = 0;
vLB('EX_ser-L[e]') = 0;
vLB('EX_spmd[e]') = 0;
vLB('EX_sucr[e]') = 0;
vLB('EX_tartr-L[e]') = 0;
vLB('EX_taur[e]') = 0;
vLB('EX_thm[e]') = 0;
vLB('EX_thr-L[e]') = 0;
vLB('EX_thymd[e]') = 0;
vLB('EX_tma[e]') = 0;
vLB('EX_tmao[e]') = 0;
vLB('EX_tre[e]') = 0;
vLB('EX_trp-L[e]') = 0;
vLB('EX_tsul[e]') = 0;
vLB('EX_ttdca[e]') = 0;
vLB('EX_tyr-L[e]') = 0;
vLB('EX_ura[e]') = 0;
vLB('EX_urea[e]') = 0;
vLB('EX_uri[e]') = 0;
vLB('EX_val-L[e]') = 0;
vLB('EX_xan[e]') = 0;
vLB('EX_xtsn[e]') = 0;
vLB('EX_xyl-D[e]') = 0;
vLB('EX_co2[e]') = 0;
vLB('EX_fe2[e]') = 0;
vLB('EX_h[e]') = 0;
vLB('EX_h2o[e]') = 0;
vLB('EX_k[e]') = 0;
vLB('EX_na1[e]') = 0;
vLB('EX_nh4[e]') = 0;
vLB('EX_pi[e]') = 0;
vLB('EX_so4[e]') = 0;

*now consider variable bounds
*aerobic conditions
vUB('EX_ac[e]') = 0;
vUB('EX_akg[e]') = 0;
vUB('EX_glc[e]') = 0;
vUB('EX_glyc[e]') = 0;
vUB('EX_lac-D[e]') = 0;
vUB('EX_lac-L[e]') = 0;
vUB('EX_mal-L[e]') = 0;
vUB('EX_o2[e]') = 0;
vUB('EX_pyr[e]') = 0;
vUB('EX_succ[e]') = 0;

vLB('EX_ac[e]') = 0;
vLB('EX_akg[e]') = 0;
vLB('EX_glc[e]') = 0;
vLB('EX_glyc[e]') = 0;
vLB('EX_lac-D[e]') = 0;
vLB('EX_lac-L[e]') = 0;
vLB('EX_mal-L[e]') = 0;
vLB('EX_o2[e]') = 0;
vLB('EX_pyr[e]') = 0;
vLB('EX_succ[e]') = 0;

v.lo(j) = -1;
v.up(j) = 1;

TFP_obj..				E =e= sum(j, eta(j));
*v_j has only one sign at any given time
TFP_const_1(j)..		v(j) =l= (1 - beta(j)) * vUB(j) - epsilon * beta(j);
TFP_const_2(j)..		v(j) =l= eta(j) * vUB(j);
TFP_const_3(j)..		v(j) =g= beta(j) * vLB(j) + epsilon * eta(j);
TFP_const_4(j)..		v(j) =g= eta(j) * vLB(j);
TFP_const_5(i)..		sum(j, S(i,j)*v(j)) =e= 0;
TFP_const_6..			sum(j,eta(j)) =l= phi;
TFP_const_7..			sum(j_db, eta(j_db)) =g= 1;
TFP_const_8(j)..		alpha(j) =l= eta(j);
TFP_const_9(j)..		alpha(j) =l= (1 - beta(j));
TFP_const_10(j)..		alpha(j) =g= eta(j) + (1 - beta(j)) - 1;
*integer cut ensure each solution is unique
TIC_int_cut_1(eqncounter)..	sum(j$(TFP_soln_f(j,eqncounter) eq 1), alpha(j)) =l= sum(j, TFP_soln_f(j,eqncounter)) - gamma(eqncounter);
TIC_int_cut_2(eqncounter)..	sum(j$(TFP_soln_b(j,eqncounter) eq 1), beta(j)) =l= sum(j, TFP_soln_b(j,eqncounter)) - (1 - gamma(eqncounter));

*define the TIC finding model
MODEL TFP
/
	TFP_obj
	TFP_const_1
	TFP_const_2
	TFP_const_3
	TFP_const_4
	TFP_const_5
	TFP_const_6
	TFP_const_7
	TFP_const_8
	TFP_const_9
	TFP_const_10
	TIC_int_cut_1
	TIC_int_cut_2
/
;

*state that there is an optfile
TFP.optfile = 1;

*treat fixed variables as constants
TFP.holdfixed = 1;

*output the possible TICs that have been found
FILE RESULT /OptFill_result_iJR904_iAF1260.txt/;
PUT RESULT;
RESULT.pw=1000;

PUT "TFP PROBLEM RESULTS"//;
PUT "REACTIONS IN TICS"/;
PUT "REACTIONS PARTICIPATING IN TIC"/;
PUT "------------------------------"/;

PUTCLOSE;

TIC_count = 0;

alias(p,p1);

Num_Solns = 0;

*repeat the solution of the TIC finding problem as long as new or unique TICs can be found
LOOP(p$(not done),
	
	/*solve the TIC finding problem*/
	temp2 = timeElapsed;
	SOLVE TFP USING MIP MINIMIZING E;
	TFP_rate(p) = timeElapsed - temp2;

	/*look at the result consider what constitutes finding a TIC*/
	/*first if the model status is 1 and E < 1 we found a TIC*/
	IF(((TFP.modelstat eq 8) or (TFP.modelstat eq 1) or (TFP.modelstat eq 2)),
		
		RESULT.ap = 1;
		PUT RESULT;
		/*write the reactions which taken together form the TIC we have found*/
		TIC_count_temp = TIC_count;
		TIC_count = TIC_count_temp + 1;
		PUT "TIC NUMBER ",TIC_count/;
		PUT "Objective Value: ",E.l:0:8/;
		PUT "Number of Reactions: ",phi:0:8/;
		PUT "Reaction            Dir  Location  eta      v(j)               alpha(j)    beta(j)"/;
		PUT '----------------------------------------------------------------------------------'/;
		
		LOOP(j_m,
		
			IF((eta.l(j_m) eq 1),
			
				RESULT.lw = 20;
				put j_m.tl;
				RESULT.lw = 12;
				
				IF((alpha.l(j_m) eq 1),
	
					put "->","   M",eta.l(j_m),system.tab,system.tab,v.l(j_m):0:8," ",alpha.l(j_m),beta.l(j_m)/;
	
				ELSEIF (beta.l(j_m) eq 1),
	
					put "<-","   M",eta.l(j_m),system.tab,system.tab,v.l(j_m):0:8,alpha.l(j_m),beta.l(j_m)/;
					
				ELSE
				
					put "XX","   M",eta.l(j_m),system.tab,system.tab,v.l(j_m):0:8,alpha.l(j_m),beta.l(j_m)/;
					
				);
				
			);
			
		);
		
		LOOP(j_db,
		
			IF((eta.l(j_db) eq 1),
			
				RESULT.lw = 20;
				put j_db.tl;
				RESULT.lw = 12;
				
				IF((alpha.l(j_db) eq 1),
	
					put "->","   DB",eta.l(j_db),system.tab,system.tab,v.l(j_db):0:8," ",alpha.l(j_db),beta.l(j_db)/;
	
				ELSEIF (beta.l(j_db) eq 1),
	
					put "<-","   DB",eta.l(j_db),system.tab,system.tab,v.l(j_db):0:8,alpha.l(j_db),beta.l(j_db)/;
				
				ELSE
				
					put "XX","   DB",eta.l(j_db),system.tab,system.tab,v.l(j_db):0:8,alpha.l(j_db),beta.l(j_db)/;
					
				);

			);
		
		);
		
		eqncounter(p) = yes;
		TFP_soln(j,eqncounter(p)) = eta.l(j);
		
		/*save directional TIC solutions*/
		TFP_soln_f(j,eqncounter(p)) = alpha.l(j);
		TFP_soln_b(j,eqncounter(p)) = beta.l(j);
		
		RESULT.ap = 1;
		PUT	RESULT;
		
		PUT "model status: ",TFP.modelstat," solver status: ",TFP.solvestat/;
		PUT "solution time: ",TFP.etSolve:0:8/;
		PUT "iterations used: ",TFP.iterUsd/;
		
		temp = Num_Solns;
		Num_Solns = temp + 1;
		
		PUT //;
		
		PUTCLOSE;
		
	/*if the model status is not one then we are out of TICs for that phi*/
	ELSE
	
		RESULT.ap = 1;
		PUT RESULT;
		
		PUT "model status: ",TFP.modelstat," solver status: ",TFP.solvestat/;
		PUT "solution time: ",TFP.etSolve:0:8/;
		PUT "iterations used: ",TFP.iterUsd/;
		PUT "completed search for phi = ",phi/;
		
		PUT //;
		
		IF((phi le phi_max),
		
			phi_temp = phi;
			phi = phi_temp + 1;
		
		ELSE
		
			/*if already reached the maximal value of phi exit the loop*/
			done = 1;
		
		);
		
	);

);

*calculate biomass growth rate statistics
IF((Num_Solns > 0),

	TFP_mean = sum(eqncounter,TFP_rate(eqncounter)) / Num_Solns;
	TFP_sd = sqrt(sum(eqncounter, power((TFP_mean - TFP_rate(eqncounter)), 2)) / Num_Solns);
	TFP_max = smax(eqncounter,TFP_rate(eqncounter));
	TFP_min = smin(eqncounter,TFP_rate(eqncounter));

ELSE

	TFP_mean = 0;
	TFP_sd = 0;
	TFP_max = 0;
	TFP_min = 0;


);

*output time to finish TIC finding
RESULT.ap = 1;
PUT RESULT;

PUT "Compilation time: ",TimeComp/;
PUT "Total elapsed time: ",TimeElapsed/;
PUT "Number of TFP solutions: ",Num_solns/;
PUT "TFP rate mean: ", TFP_mean:0:8/;
PUT "TFP rate standard deviaton: ", TFP_sd:0:8/;
PUT "TFP rate max: ", TFP_max:0:8/;
PUT "TFP rate min: ", TFP_min:0:8/;
PUT "Compilation time: ",TimeComp/;
PUT "Elapsed time to find all TICs: ",TimeElapsed/;
PUT "Number of TICs identified: ",TIC_count//;

PUTCLOSE;

**************************** Connecting Problems ****************************************
*define the variables for the primary problem
BINARY VARIABLES

	delta(j_db)			value of 1 if reaction j_db is being added irreversibly forwards from the database
	rho(j_db)			value of 1 if reaction j_db is being added irreversibly backwards from the database
	omega(j_db)			value of 1 if reaction j_db is being added reversibly from the database essentially delta times rho
	sigma(p)			used to enforce integer cuts related to delta and iota ensures set of deltas or set of iotas can be the same for two solutions but not both
	tau(p)				used to enforce integer cuts related to delta and iota ensures set of deltas or set of iotas can be the same for a TIC and a solution but not both
	w(i,j)				value of 1 if reaction j produces metabolite i otherwise 0
	x(i)				value of 1 if model can produce metabolite i
	zeta(j_db)			value of 1 if reaction j_db is being added 0 otherwise
	theta(j)
	lambda(j)

VARIABLES

	v_2(j)				value of reaction rate
	met_obj				maximum number of metabolites connected in the network
	rxn_obj				minimum number of reactions added to achieve such connections
	omega_obj			maximum number of reversible reactions added to achieve max connections with minimum reactions

;

*define primary equations
EQUATIONS

	find_met_obj
	find_rxn_obj
	find_omega_obj
	CPs_const_1
	CPs_const_1b
	CPs_const_2(j_db)
	CPs_const_3(j_db)
	CPs_const_4(biomass_precursors)
	CPs_const_5(i)
	CPs_const_6(j_db)
	CPs_const_7(j_db)
	CPs_const_8(j_db)
	CPs_const_9(j)
	CPs_const_10(j)
	CPs_const_11(j)
	CPs_const_12(j)
	CPs_const_13(i)
	CPs_const_14(j_db)
	int_cut_1(p)
	int_cut_2(p)
	int_cut_3(p)
	int_cut_4(p)
	int_cut_5(p)
	rxn_const_1
	omega_const_1
	

PARAMETERS

	max_connections
	min_rxns

;

*initialize max connections
max_connections = 0;

find_met_obj..						met_obj =e= sum(i_m, x(i_m));
CPs_const_1..						sum(j_db, zeta(j_db)) =g= 1;
CPs_const_1b..						sum(j_db, zeta(j_db)) =e= 0;
CPs_const_2(j_db)..					v_2(j_db) =l= delta(j_db) * vUB(j_db);
CPs_const_3(j_db)..					v_2(j_db) =g= rho(j_db) * vLB(j_db);
CPs_const_4(biomass_precursors)..	x(biomass_precursors) =e= 1;
CPs_const_5(i)..					sum(j, S(i,j) * v_2(j)) =e= 0;
CPs_const_6(j_db)..					delta(j_db) + rho(j_db) - omega(j_db) =e= zeta(j_db);
CPs_const_7(j_db)..					omega(j_db) =l= delta(j_db);
CPs_const_8(j_db)..					omega(j_db) =l= rho(j_db);
CPs_const_9(j)..					v_2(j) =l= (1 - theta(j)) * vUB(j) - epsilon * theta(j);
CPs_const_10(j)..					v_2(j) =g= theta(j) * vLB(j);
CPs_const_11(j)..					v_2(j) =l= lambda(j) * vUB(j);
CPs_const_12(j)..					v_2(j) =g= (1-lambda(j)) * vLB(j) + epsilon * lambda(j);
CPs_const_13(i)..					x(i) =l= sum(j,lambda(j) * xi(i,j) + theta(j) * psi(i,j));
CPs_const_14(j_db)..				zeta(j_db) =l= sum(i,lambda(j_db) * xi(i,j_db) + theta(j_db) * psi(i,j_db));
int_cut_1(eqncounter_2)..			sum(j_db$(CPs_soln_f(j_db,eqncounter_2) eq 1), delta(j_db)) =l= sum(j_db, CPs_soln_f(j_db,eqncounter_2)) - sigma(eqncounter_2);
int_cut_2(eqncounter_2)..			sum(j_db$(CPs_soln_b(j_db,eqncounter_2) eq 1), rho(j_db)) =l= sum(j_db, CPs_soln_b(j_db,eqncounter_2)) - (1 - sigma(eqncounter_2));
int_cut_3(eqncounter_2)..			sum(j_db$(CPs_soln_f(j_db,eqncounter_2) eq 1), CPs_soln_f(j_db,eqncounter_2) - delta(j_db)) + sum(j_db$(CPs_soln_b(j_db,eqncounter_2) eq 1), CPs_soln_b(j_db,eqncounter_2) - rho(j_db)) =g= num_omegas(eqncounter_2) + 1;
int_cut_4(eqncounter)..				sum(j_db$(TFP_soln_f(j_db,eqncounter) eq 1), delta(j_db)) =l= sum(j_db, TFP_soln_f(j_db,eqncounter)) - tau(eqncounter);
int_cut_5(eqncounter)..				sum(j_db$(TFP_soln_b(j_db,eqncounter) eq 1), rho(j_db)) =l= sum(j_db, TFP_soln_b(j_db,eqncounter)) - (1 - tau(eqncounter));

*define the primary model
MODEL CP1
/
	find_met_obj
	CPs_const_1
	CPs_const_2
	CPs_const_3
	CPs_const_4
	CPs_const_5
	CPs_const_6
	CPs_const_7
	CPs_const_8
	int_cut_1
	int_cut_2
	int_cut_3
	int_cut_4
	int_cut_5
	CPs_const_9
	CPs_const_10
	CPs_const_11
	CPs_const_12
	CPs_const_13
	CPs_const_14
/
;

*model to solve for minimum number of reactions to reach the number of connections
find_rxn_obj..	rxn_obj =e= sum(j_db, zeta(j_db));
rxn_const_1..	sum(i_m,x(i_m)) =e= max_connections;

MODEL CP2
/
	find_rxn_obj
	rxn_const_1
	CPs_const_1
	CPs_const_2
	CPs_const_3
	CPs_const_4
	CPs_const_5
	CPs_const_6
	CPs_const_7
	CPs_const_8
	int_cut_1
	int_cut_2
	int_cut_3
	int_cut_4
	int_cut_5
	CPs_const_9
	CPs_const_10
	CPs_const_11
	CPs_const_12
	CPs_const_13
	CPs_const_14
	
/
;

*model to solve for the maximum number of omega values e.g. only add irreversible reactions when necessary
find_omega_obj..	omega_obj =e= sum(j_db, omega(j_db));
omega_const_1..		sum(j_db, zeta(j_db)) =e= min_rxns;

MODEL CP3
/
	find_omega_obj
	rxn_const_1
	omega_const_1
	CPs_const_1
	CPs_const_2
	CPs_const_3
	CPs_const_4
	CPs_const_5
	CPs_const_6
	CPs_const_7
	CPs_const_8
	int_cut_1
	int_cut_2
	int_cut_3
	int_cut_4
	int_cut_5
	CPs_const_9
	CPs_const_10
	CPs_const_11
	CPs_const_12
	CPs_const_13
	CPs_const_14
/
;

MODEL GAPFIND
/
	find_met_obj
	CPs_const_1b
	CPs_const_2
	CPs_const_3
	CPs_const_4
	CPs_const_5
	CPs_const_6
	CPs_const_7
	CPs_const_8
	int_cut_1
	int_cut_2
	int_cut_3
	int_cut_4
	int_cut_5
	CPs_const_9
	CPs_const_10
	CPs_const_11
	CPs_const_12
	CPs_const_13
	CPs_const_14
/
;

done = 0;
alias(p1,p);

*add in layer to perform FBA with maximizing biomass to pull some metrics about model differences
*resulting from different solutions. 
EQUATIONS

	growth_obj
	mass_bal(i)
	UB_m(j_m)
	UB_db(j_db)
	LB_m(j_m)
	LB_db(j_db)

;

VARIABLES

	Z			growth objective variable
	v_3(j)		value of reaction rate in FBA

;

*require ATPM to be met for connecting solution and FBA analysis
vUB('ATPM') = 2;
vLB('ATPM') = 2;

*turn on growth medium for connecting problems
*first consider constant bounds
vUB('EX_12ppd-S[e]') = M;
vUB('EX_15dap[e]') = M;
vUB('EX_26dap-M[e]') = M;
vUB('EX_2ddglcn[e]') = M;
vUB('EX_3hcinnm[e]') = M;
vUB('EX_3hpppn[e]') = M;
vUB('EX_4abut[e]') = M;
vUB('EX_acac[e]') = M;
vUB('EX_acald[e]') = M;
vUB('EX_acgam[e]') = M;
vUB('EX_acmana[e]') = M;
vUB('EX_acnam[e]') = M;
vUB('EX_ade[e]') = M;
vUB('EX_adn[e]') = M;
vUB('EX_ala-D[e]') = M;
vUB('EX_ala-L[e]') = M;
vUB('EX_alltn[e]') = M;
vUB('EX_amp[e]') = M;
vUB('EX_arab-L[e]') = M;
vUB('EX_arg-L[e]') = M;
vUB('EX_asn-L[e]') = M;
vUB('EX_asp-L[e]') = M;
vUB('EX_but[e]') = M;
vUB('EX_cbl1[e]') = M;
vUB('EX_chol[e]') = M;
vUB('EX_cit[e]') = M;
vUB('EX_crn[e]') = M;
vUB('EX_csn[e]') = M;
vUB('EX_cynt[e]') = M;
vUB('EX_cys-L[e]') = M;
vUB('EX_cytd[e]') = M;
vUB('EX_dad-2[e]') = M;
vUB('EX_dcyt[e]') = M;
vUB('EX_dgsn[e]') = M;
vUB('EX_dha[e]') = M;
vUB('EX_din[e]') = M;
vUB('EX_dms[e]') = M;
vUB('EX_dmso[e]') = M;
vUB('EX_duri[e]') = M;
vUB('EX_etoh[e]') = M;
vUB('EX_for[e]') = M;
vUB('EX_fru[e]') = M;
vUB('EX_fuc-L[e]') = M;
vUB('EX_fuc1p-L[e]') = M;
vUB('EX_fum[e]') = M;
vUB('EX_g6p[e]') = M;
vUB('EX_gal[e]') = M;
vUB('EX_galct-D[e]') = M;
vUB('EX_galctn-D[e]') = M;
vUB('EX_galt[e]') = M;
vUB('EX_galur[e]') = M;
vUB('EX_gam[e]') = M;
vUB('EX_gbbtn[e]') = M;
vUB('EX_glcn[e]') = M;
vUB('EX_glcr[e]') = M;
vUB('EX_glcur[e]') = M;
vUB('EX_gln-L[e]') = M;
vUB('EX_glu-L[e]') = M;
vUB('EX_gly[e]') = M;
vUB('EX_glyald[e]') = M;
vUB('EX_glyb[e]') = M;
vUB('EX_glyc3p[e]') = M;
vUB('EX_glyclt[e]') = M;
vUB('EX_gsn[e]') = M;
vUB('EX_gua[e]') = M;
vUB('EX_hdca[e]') = M;
vUB('EX_his-L[e]') = M;
vUB('EX_hxan[e]') = M;
vUB('EX_idon-L[e]') = M;
vUB('EX_ile-L[e]') = M;
vUB('EX_indole[e]') = M;
vUB('EX_ins[e]') = M;
vUB('EX_lcts[e]') = M;
vUB('EX_leu-L[e]') = M;
vUB('EX_lys-L[e]') = M;
vUB('EX_malt[e]') = M;
vUB('EX_malthx[e]') = M;
vUB('EX_maltpt[e]') = M;
vUB('EX_malttr[e]') = M;
vUB('EX_maltttr[e]') = M;
vUB('EX_man[e]') = M;
vUB('EX_man6p[e]') = M;
vUB('EX_melib[e]') = M;
vUB('EX_met-D[e]') = M;
vUB('EX_met-L[e]') = M;
vUB('EX_mnl[e]') = M;
vUB('EX_nac[e]') = M;
vUB('EX_nad[e]') = M;
vUB('EX_nmn[e]') = M;
vUB('EX_no2[e]') = M;
vUB('EX_no3[e]') = M;
vUB('EX_ocdca[e]') = M;
vUB('EX_orn[e]') = M;
vUB('EX_phe-L[e]') = M;
vUB('EX_pnto-R[e]') = M;
vUB('EX_pppn[e]') = M;
vUB('EX_pro-L[e]') = M;
vUB('EX_ptrc[e]') = M;
vUB('EX_rib-D[e]') = M;
vUB('EX_rmn[e]') = M;
vUB('EX_sbt-D[e]') = M;
vUB('EX_ser-D[e]') = M;
vUB('EX_ser-L[e]') = M;
vUB('EX_spmd[e]') = M;
vUB('EX_sucr[e]') = M;
vUB('EX_tartr-L[e]') = M;
vUB('EX_taur[e]') = M;
vUB('EX_thm[e]') = M;
vUB('EX_thr-L[e]') = M;
vUB('EX_thymd[e]') = M;
vUB('EX_tma[e]') = M;
vUB('EX_tmao[e]') = M;
vUB('EX_tre[e]') = M;
vUB('EX_trp-L[e]') = M;
vUB('EX_tsul[e]') = M;
vUB('EX_ttdca[e]') = M;
vUB('EX_tyr-L[e]') = M;
vUB('EX_ura[e]') = M;
vUB('EX_urea[e]') = M;
vUB('EX_uri[e]') = M;
vUB('EX_val-L[e]') = M;
vUB('EX_xan[e]') = M;
vUB('EX_xtsn[e]') = M;
vUB('EX_xyl-D[e]') = M;
vUB('EX_co2[e]') = M;
vUB('EX_fe2[e]') = M;
vUB('EX_h[e]') = M;
vUB('EX_h2o[e]') = M;
vUB('EX_k[e]') = M;
vUB('EX_na1[e]') = M;
vUB('EX_nh4[e]') = M;
vUB('EX_pi[e]') = M;
vUB('EX_so4[e]') = M;

vLB('EX_12ppd-S[e]') = 0;
vLB('EX_15dap[e]') = 0;
vLB('EX_26dap-M[e]') = 0;
vLB('EX_2ddglcn[e]') = 0;
vLB('EX_3hcinnm[e]') = 0;
vLB('EX_3hpppn[e]') = 0;
vLB('EX_4abut[e]') = 0;
vLB('EX_acac[e]') = 0;
vLB('EX_acald[e]') = 0;
vLB('EX_acgam[e]') = 0;
vLB('EX_acmana[e]') = 0;
vLB('EX_acnam[e]') = 0;
vLB('EX_ade[e]') = 0;
vLB('EX_adn[e]') = 0;
vLB('EX_ala-D[e]') = 0;
vLB('EX_ala-L[e]') = 0;
vLB('EX_alltn[e]') = 0;
vLB('EX_amp[e]') = 0;
vLB('EX_arab-L[e]') = 0;
vLB('EX_arg-L[e]') = 0;
vLB('EX_asn-L[e]') = 0;
vLB('EX_asp-L[e]') = 0;
vLB('EX_but[e]') = 0;
vLB('EX_cbl1[e]') = 0;
vLB('EX_chol[e]') = 0;
vLB('EX_cit[e]') = 0;
vLB('EX_crn[e]') = 0;
vLB('EX_csn[e]') = 0;
vLB('EX_cynt[e]') = 0;
vLB('EX_cys-L[e]') = 0;
vLB('EX_cytd[e]') = 0;
vLB('EX_dad-2[e]') = 0;
vLB('EX_dcyt[e]') = 0;
vLB('EX_dgsn[e]') = 0;
vLB('EX_dha[e]') = 0;
vLB('EX_din[e]') = 0;
vLB('EX_dms[e]') = 0;
vLB('EX_dmso[e]') = 0;
vLB('EX_duri[e]') = 0;
vLB('EX_etoh[e]') = 0;
vLB('EX_for[e]') = 0;
vLB('EX_fru[e]') = 0;
vLB('EX_fuc-L[e]') = 0;
vLB('EX_fuc1p-L[e]') = 0;
vLB('EX_fum[e]') = 0;
vLB('EX_g6p[e]') = 0;
vLB('EX_gal[e]') = 0;
vLB('EX_galct-D[e]') = 0;
vLB('EX_galctn-D[e]') = 0;
vLB('EX_galt[e]') = 0;
vLB('EX_galur[e]') = 0;
vLB('EX_gam[e]') = 0;
vLB('EX_gbbtn[e]') = 0;
vLB('EX_glcn[e]') = 0;
vLB('EX_glcr[e]') = 0;
vLB('EX_glcur[e]') = 0;
vLB('EX_gln-L[e]') = 0;
vLB('EX_glu-L[e]') = 0;
vLB('EX_gly[e]') = 0;
vLB('EX_glyald[e]') = 0;
vLB('EX_glyb[e]') = 0;
vLB('EX_glyc3p[e]') = 0;
vLB('EX_glyclt[e]') = 0;
vLB('EX_gsn[e]') = 0;
vLB('EX_gua[e]') = 0;
vLB('EX_hdca[e]') = 0;
vLB('EX_his-L[e]') = 0;
vLB('EX_hxan[e]') = 0;
vLB('EX_idon-L[e]') = 0;
vLB('EX_ile-L[e]') = 0;
vLB('EX_indole[e]') = 0;
vLB('EX_ins[e]') = 0;
vLB('EX_lcts[e]') = 0;
vLB('EX_leu-L[e]') = 0;
vLB('EX_lys-L[e]') = 0;
vLB('EX_malt[e]') = 0;
vLB('EX_malthx[e]') = 0;
vLB('EX_maltpt[e]') = 0;
vLB('EX_malttr[e]') = 0;
vLB('EX_maltttr[e]') = 0;
vLB('EX_man[e]') = 0;
vLB('EX_man6p[e]') = 0;
vLB('EX_melib[e]') = 0;
vLB('EX_met-D[e]') = 0;
vLB('EX_met-L[e]') = 0;
vLB('EX_mnl[e]') = 0;
vLB('EX_nac[e]') = 0;
vLB('EX_nad[e]') = 0;
vLB('EX_nmn[e]') = 0;
vLB('EX_no2[e]') = 0;
vLB('EX_no3[e]') = 0;
vLB('EX_ocdca[e]') = 0;
vLB('EX_orn[e]') = 0;
vLB('EX_phe-L[e]') = 0;
vLB('EX_pnto-R[e]') = 0;
vLB('EX_pppn[e]') = 0;
vLB('EX_pro-L[e]') = 0;
vLB('EX_ptrc[e]') = 0;
vLB('EX_rib-D[e]') = 0;
vLB('EX_rmn[e]') = 0;
vLB('EX_sbt-D[e]') = 0;
vLB('EX_ser-D[e]') = 0;
vLB('EX_ser-L[e]') = 0;
vLB('EX_spmd[e]') = 0;
vLB('EX_sucr[e]') = 0;
vLB('EX_tartr-L[e]') = 0;
vLB('EX_taur[e]') = 0;
vLB('EX_thm[e]') = 0;
vLB('EX_thr-L[e]') = 0;
vLB('EX_thymd[e]') = 0;
vLB('EX_tma[e]') = 0;
vLB('EX_tmao[e]') = 0;
vLB('EX_tre[e]') = 0;
vLB('EX_trp-L[e]') = 0;
vLB('EX_tsul[e]') = 0;
vLB('EX_ttdca[e]') = 0;
vLB('EX_tyr-L[e]') = 0;
vLB('EX_ura[e]') = 0;
vLB('EX_urea[e]') = 0;
vLB('EX_uri[e]') = 0;
vLB('EX_val-L[e]') = 0;
vLB('EX_xan[e]') = 0;
vLB('EX_xtsn[e]') = 0;
vLB('EX_xyl-D[e]') = 0;
vLB('EX_co2[e]') = -M;
vLB('EX_fe2[e]') = -M;
vLB('EX_h[e]') = -M;
vLB('EX_h2o[e]') = -M;
vLB('EX_k[e]') = -M;
vLB('EX_na1[e]') = -M;
vLB('EX_nh4[e]') = -M;
vLB('EX_pi[e]') = -M;
vLB('EX_so4[e]') = -M;

*now consider variable bounds
*aerobic conditions
vUB('EX_ac[e]') = M;
vUB('EX_akg[e]') = M;
vUB('EX_glc[e]') = M;
vUB('EX_glyc[e]') = M;
vUB('EX_lac-D[e]') = M;
vUB('EX_lac-L[e]') = M;
vUB('EX_mal-L[e]') = M;
vUB('EX_o2[e]') = 0;
vUB('EX_pyr[e]') = M;
vUB('EX_succ[e]') = M;

vLB('EX_ac[e]') = -5;
vLB('EX_akg[e]') = 0;
vLB('EX_glc[e]') = 0;
vLB('EX_glyc[e]') = 0;
vLB('EX_lac-D[e]') = 0;
vLB('EX_lac-L[e]') = 0;
vLB('EX_mal-L[e]') = 0;
vLB('EX_o2[e]') = -20;
vLB('EX_pyr[e]') = 0;
vLB('EX_succ[e]') = 0;

v_2.lo(j) = vLB(j);
v_2.up(j) = vUB(j);
v_3.lo(j) = vLB(j);
v_3.up(j) = vUB(j);

growth_obj..	Z =e= v_3('biomass');
mass_bal(i)..	sum(j, S(i,j) * v_3(j)) =e= 0;
UB_m(j_m)..		v_3(j_m) =l= vUB(j_m);
UB_db(j_db)..	v_3(j_db) =l= curr_CPs_soln_f(j_db) * vUB(j_db);
LB_m(j_m)..		v_3(j_m) =g= vLB(j_m);
LB_db(j_db)..	v_3(j_db) =g= curr_CPs_soln_b(j_db) * vLB(j_db);

MODEL FBA
/
	growth_obj
	mass_bal
	UB_m
	UB_db
	LB_m
	LB_db
/
;

*state that there is an optfile
FBA.optfile = 1;
CP1.optfile = 6;
CP2.optfile = 6;
CP3.optfile = 6;

Num_Solns = 0;

*solve a base-case to see the maximum number of metaboltes which can be produced
SOLVE GAPFIND USING MIP MAXIMIZING met_obj;

PUT "Number of metabolites produced by raw model: ", met_obj.l/;
base_prod(i_m) = x.l(i_m);

PUT //;

LOOP(p$(not done),
	
	/*need to build in some sort of error handling mechanism*/
	iter = 0;
	iter_max = 3;
	
	/*find the maximum number of reactions which can be fit*/
	WHILE(((iter = 0) OR ((iter le iter_max) AND (CP1.modelStat ne 1))),
		
		/*if this is the second solution or later relax the feasibility constraints*/
		IF((iter ge 1),
			
			CP1.optfile = 7;
			
		);
			
		/*now that we know the maximum connections minimize the number of metabolites*/
		/*which can be fixed now find the minimum number of reactions corresponding*/
		/*to that*/
		/*put framework around this statement to get the solution time*/
		temp2 = timeElapsed;
		SOLVE CP1 USING MIP MAXIMIZING met_obj;
		time_cp1 = timeElapsed - temp2;
		max_connections = met_obj.l;
			
		temp = iter;
		iter = temp + 1;
		
	);
	
	iter_CP1 = iter;

	IF((CP1.modelstat eq 1),
		
		/*need to build in some sort of error handling mechanism*/
		iter = 0;
		iter_max = 5;
		
		WHILE(((iter = 0) OR ((iter le iter_max) AND (CP2.modelStat ne 1))),
		
			/*if this is the second solution or later relax the feasibility constraints*/
			IF((iter ge 1),
			
				CP2.optfile = 7;
			
			);
			
			/*now that we know the maximum connections minimize the number of metabolites*/
			/*which can be fixed now find the minimum number of reactions corresponding*/
			/*to that*/
			/*put framework around this statement to get the solution time*/
			temp2 = timeElapsed;
			SOLVE CP2 USING MIP MINIMIZING rxn_obj;
			time_cp2 = timeElapsed - temp2;
			min_rxns = rxn_obj.l;
			
			temp = iter;
			iter = temp + 1;
		
		);
		
		iter_CP2 = iter;
		
		/*reset problem constraints on feasibility*/
		CP2.optfile = 6;
		
		/*ideally will have an optimal or at least feasible solution by this point*/
		
		/*same structure of error handling for second inner problem*/
		iter = 0;
		iter_max = 5;
		
		WHILE(((iter = 0) OR ((iter le iter_max) AND (CP3.modelStat ne 1))),
		
			/*if this is the second solution or later relax the feasibility constraints*/
			IF((iter ge 1),
			
				CP3.optfile = 7;
			
			);
			
			/*now that we know the miminum number of reactions which we can add to achieve the*/
			/*maximum connectivity make sure we add as many reactions reversibly as possible*/
			/*put framework around this statement to get the solution time*/
			temp2 = timeElapsed;
			SOLVE CP3 USING MIP MAXIMIZING omega_obj;
			time_cp3 = timeElapsed - temp2;
			
			temp = iter;
			iter = temp + 1;
		
		);
		
		iter_CP3 = iter;
		
		CP_rate(p) = time_cp1 + time_cp2 + time_cp3;
		
		/*reset problem constraints on feasibility*/
		CP3.optfile = 6;
	
		RESULT.ap = 1;
		PUT RESULT;

		/*write the solution results to the output file*/
		PUT "************************           CONNECTING PROBLEMS SOLUTION ",p.tl,"************************"//;
		PUT "OUTER OBJECTIVE VALUE: ",max_connections/;
		PUT "INNER OBJECTIVE VALUE 1: ",min_rxns/;
		PUT "INNER OBJECTIVE VALUE 2: ",omega_obj.l//;
		PUT "RXN TO ADD           DIRECTION   DELTA       RHO         OMEGA        ZETA  SOLVED RATE"/;
		PUT "------------------------------------------------------------------------------------------"/;

		count_rxns = 0;

		LOOP(j_db,

			IF(((omega.l(j_db) le (1 + epsilon)) AND (omega.l(j_db) ge (1 - epsilon))),
	
				RESULT.lw = 20;
				PUT j_db.tl;
				RESULT.lw = 12;
			
				count_rxns_temp = count_rxns;
				count_rxns = count_rxns_temp + 1;
	
				PUT " <-> ",delta.l(j_db),rho.l(j_db),omega.l(j_db),zeta.l(j_db),system.tab,v_2.l(j_db):0:8/;
			
			ELSEIF ((delta.l(j_db) le (1 + epsilon)) AND (delta.l(j_db) ge (1 - epsilon))),
	
				RESULT.lw = 20;
				PUT j_db.tl;
				RESULT.lw = 12;
			
				count_rxns_temp = count_rxns;
				count_rxns = count_rxns_temp + 1;
	
				PUT " ->  ",delta.l(j_db),rho.l(j_db),omega.l(j_db),zeta.l(j_db),system.tab,v_2.l(j_db):0:8/;
			
			ELSEIF ((rho.l(j_db) le (1 + epsilon)) AND (rho.l(j_db) ge (1 - epsilon))),
	
				RESULT.lw = 20;
				PUT j_db.tl;
				RESULT.lw = 12;
			
				count_rxns_temp = count_rxns;
				count_rxns = count_rxns_temp + 1;
	
				PUT " <-  ",delta.l(j_db),rho.l(j_db),omega.l(j_db),zeta.l(j_db),system.tab,v_2.l(j_db):0:8/;
				
			ELSE
			
				RESULT.lw = 20;
				PUT j_db.tl;
				RESULT.lw = 12;
				
				PUT " XX  ",delta.l(j_db),rho.l(j_db),omega.l(j_db),zeta.l(j_db),system.tab,v_2.l(j_db):0:8/;
				
			);

		);

		PUT "Total reactions to be added: ",count_rxns, " (should be: ",min_rxns,")";
		
		PUT //;
		
		PUT "MODEL RXN            DIRECTION  THETA        LAMBDA     SOLVED RATE"/;
		PUT "--------------------------------------------------------------------"/;

		count_rxns = 0;

		LOOP(j_m,

			IF (((theta.l(j_m) le (1 + epsilon)) AND (theta.l(j_m) ge (1 - epsilon))),
	
				RESULT.lw = 20;
				PUT j_m.tl;
				RESULT.lw = 12;
	
				PUT " <-  ",theta.l(j_m),lambda.l(j_m),system.tab,system.tab,v_2.l(j_m):0:8/;
				
			ELSEIF ((lambda.l(j_m) le (1 + epsilon)) AND (lambda.l(j_m) ge (1 - epsilon))),
	
				RESULT.lw = 20;
				PUT j_m.tl;
				RESULT.lw = 12;
	
				PUT " ->  ",theta.l(j_m),lambda.l(j_m),system.tab,system.tab,v_2.l(j_m):0:8/;

			ELSE
			
				RESULT.lw = 20;
				PUT j_m.tl;
				RESULT.lw = 12;
	
				PUT " XX  ",theta.l(j_m),lambda.l(j_m),system.tab,system.tab,v_2.l(j_m):0:8/;
			);

		);
		
		PUT //;
		
		PUT "METABOLITES NEWLY PRODUCABLE THROUGH OPTFILL"/;
		PUT '--------------------------------------------'/;
		
		LOOP(i_m, 
		
			IF(((x.l(i_m) eq 1) AND (x.l(i_m) ne base_prod(i_m))),
			
				RESULT.lw = 20;
				PUT i_m.tl/; 
				RESULT.lw = 12; 
			
			);
			
		);

		PUT //;

		PUT 'METABOLITES THAT CAN NOW BE PRODUCED'/;
		PUT 'Metabolite  ','Origin   ','x(i)'/;
		PUT '------------------------------------'/;

		LOOP(i_m,

			RESULT.lw = 20;
			PUT i_m.tl;
			RESULT.lw = 12;
			
			
			PUT ' M ',x.l(i_m)/;

		);

		LOOP(i_db,

			RESULT.lw = 20;
			PUT i_db.tl;
			RESULT.lw = 12;
			
			PUT ' DB',x.l(i_db)/;

		);
		
		PUT //;
		
		curr_CPs_soln_f(j_db) = delta.l(j_db);
		curr_CPs_soln_b(j_db) = rho.l(j_db);
		
		/*solve FBA to get some characteristics of the current connecting solution*/
		SOLVE FBA USING LP MAXIMIZING Z;
		
		PUT 'FLUX RATES OF THE CURRENT SOLUTION (FBA, max growth)'/;
		PUT 'Reaction    Origin   rate               UB          UB          LB          LB'/;
		PUT '--------------------------------------------------------------------------------'/;

		LOOP(j_m,

			RESULT.lw = 20;
			PUT j_m.tl;
			RESULT.lw = 12; 
			
			PUT '  M    ',system.tab,v_3.l(j_m):0:8,system.tab,v_3.up(j_m),vUB(j_m),v_3.lo(j_m),vLB(j_m)/;

		);

		LOOP(j_db,
		
			IF(((omega.l(j_db) le (1 + epsilon)) AND ((omega.l(j_db) ge (1 - epsilon)))),
			
				RESULT.lw = 20;
				PUT j_db.tl;
				RESULT.lw = 12; 
				
				PUT '  DB<->',system.tab,system.tab,v_3.l(j_db):0:8,system.tab,v_3.up(j_db),(curr_CPs_soln_f(j_db) * vUB(j_db)),v_3.lo(j_db),(curr_CPs_soln_b(j_db) * vLB(j_db))/;
			
			ELSEIF ((delta.l(j_db) le (1 + epsilon)) AND (delta.l(j_db) ge (1 - epsilon))),
			
				RESULT.lw = 20;
				PUT j_db.tl;
				RESULT.lw = 12; 
				
				PUT '  DB-> ',system.tab,system.tab,v_3.l(j_db):0:8,system.tab,v_3.up(j_db),vUB(j_db),v_3.lo(j_db),vLB(j_db)/;
			
			ELSEIF ((rho.l(j_db) le (1 + epsilon)) AND (rho.l(j_db) ge (1 - epsilon))),
			
				RESULT.lw = 20;
				PUT j_db.tl;
				RESULT.lw = 12; 
				
				PUT '  DB<- ',system.tab,system.tab,v_3.l(j_db):0:8,system.tab,v_3.up(j_db),vUB(j_db),v_3.lo(j_db),vLB(j_db)/;
			
			);

		);
		
		PUT "CP1: Model status: ",CP1.modelstat," solver status: ",CP1.solvestat," (iterations ",iter_CP1,")"/;
		PUT "CP2: Model status: ",CP2.modelstat," solver status: ",CP2.solvestat," (iterations ",iter_CP2,")"/;
		PUT "CP3: Model status: ",CP3.modelstat," solver status: ",CP3.solvestat," (iterations ",iter_CP3,")"/;
		PUT "FBA: Model status: ",FBA.modelstat," solver status: ",FBA.solvestat/;
		
		/*add this to current list of solutions*/
		eqncounter_2(p)=yes;
		CPs_soln(j_db,eqncounter_2(p))=delta.l(j_db) + rho.l(j_db) - omega.l(j_db);
		CPs_soln_f(j_db,eqncounter_2(p))=delta.l(j_db);
		CPs_soln_b(j_db,eqncounter_2(p))=rho.l(j_db);
		num_omegas(eqncounter_2(p))=sum(j_db, omega.l(j_db));
		temp = sum(j_db, omega.l(j_db));
		PUT "Number omegas: ",temp/;
		
		done = 0;
		
		temp = Num_Solns;
		Num_Solns = temp + 1;
		
		/*report the results of the FBA*/
		PUT "new model biomass rate: ", Z.l:0:8/;
		PUT "solve time: ", CP_rate(p):0:8/;
		
		bio_rate(p) = Z.l;
		
		PUT //;

		PUTCLOSE;
		
	ELSE
	
		/*otherwise there are no more solutions left*/
		
		RESULT.ap = 1;
		PUT RESULT;

		/*write the solution results to the output file*/
		PUT "************************           END OF SOLUTIONS           ************************"//;
		PUT "No solution found, model status: ",CP1.modelstat," solver status: ",CP1.solvestat/;
		
		PUTCLOSE;
		
		done = 1;
		
	);
	
);

*calculate biomass growth rate statistics
IF((Num_Solns > 0),

	bio_mean = sum(eqncounter_2,bio_rate(eqncounter_2)) / Num_Solns;
	bio_sd = sqrt(sum(eqncounter_2, power((bio_mean - bio_rate(eqncounter_2)), 2)) / Num_Solns);
	bio_max = smax(eqncounter_2,bio_rate(eqncounter_2));
	bio_min = smin(eqncounter_2,bio_rate(eqncounter_2));

ELSE

	bio_mean = 0;
	bio_sd = 0;
	bio_max = 0;
	bio_min = 0;


);

*calculate biomass growth rate statistics
IF((Num_Solns > 0),

	CP_mean = sum(eqncounter_2,CP_rate(eqncounter_2)) / Num_Solns;
	CP_sd = sqrt(sum(eqncounter_2, power((CP_mean - CP_rate(eqncounter_2)), 2)) / Num_Solns);
	CP_max = smax(eqncounter_2,CP_rate(eqncounter_2));
	CP_min = smin(eqncounter_2,CP_rate(eqncounter_2));

ELSE

	CP_mean = 0;
	CP_sd = 0;
	CP_max = 0;
	CP_min = 0;


);

*output time to finish TIC finding
RESULT.ap = 1;
PUT RESULT;
PUT "Compilation time: ",TimeComp/;
PUT "Total elapsed time: ",TimeElapsed/;
PUT "Number of connecting solutions: ",Num_solns/;
PUT "Biomass rate mean: ", bio_mean:0:8/;
PUT "Biomass rate standard deviaton: ", bio_sd:0:8/;
PUT "Biomass rate max: ", bio_max:0:8/;
PUT "Biomass rate min: ", bio_min:0:8/;
PUT "CP rate mean: ", CP_mean:0:8/;
PUT "CP rate standard deviaton: ", CP_sd:0:8/;
PUT "CP rate max: ", CP_max:0:8/;
PUT "CP rate min: ", CP_min:0:8//;
PUTCLOSE;
```
