## Supplemental File 3 for "A Novel Optimization-Based Tool to Automate Infeasible Cycle-Free Gapfilling of Genome-Scale Metabolic Models": prep_for_optfill_iJR904_iAF1260.pl

```
#!usr/bin/perl -w

#Written by: Wheaton Schroeder
#Written to prepare all the datafiles necessary to do OptFill on the ED model

require "perl_lib_1.pm";
use strict;

system("python convertWLS_model.py");
system("python convertWLS_db.py");

printf "\n\nSuccessfully ran convert scripts\n\n";

#get model reaction list
open(MODELRXNS, "<rxns_m.txt") or die "Could not open the model reaction list file, reason: $!\n";
chomp(my @model_rxns = <MODELRXNS>);
#remove the "/" characters
pop @model_rxns;
shift @model_rxns;

#get the model metabolite list
open(MODELMETS, "<mets_m.txt") or die "Could not open the model metabolite list file, reason: $!\n";
chomp(my @model_mets = <MODELMETS>);
pop @model_mets;
shift @model_mets;

#get the database reaction list
open(DBMETS, "<rxns_db.txt") or die "Could not open the database reaction list file, reason: $!\n";
chomp(my @db_rxns = <DBMETS>);
pop @db_rxns;
shift @db_rxns;

#get the database metabolite list
open(DBMETS, "<mets_db.txt") or die "Could not open the database metabolite list file, reason: $!\n";
chomp(my @db_mets = <DBMETS>);
pop @db_mets;
shift @db_mets;

#create the all_mets file
open(ALLMETS, ">all_mets.txt") or die "could not write to all_mets.txt, reason: $!\n";

#write the leading "/"
printf ALLMETS "\/\n";

#combine the two metabolite lists
my @all_mets = @db_mets;

for(my $a=0; $a <= $#model_mets; $a++) {

	push @all_mets, $model_mets[$a];

}  

#remove redundant metabolites
my $nr_all_mets_ref = &RemoveRedundancies(\@all_mets);
my @nr_all_mets = @$nr_all_mets_ref;

#write the non-redundante metabolite list
for(my $b=0; $b <= $#nr_all_mets; $b++) {

	printf ALLMETS "%s\n", $nr_all_mets[$b];

}

#put trailing "/"
printf ALLMETS "\/";

#write the all_rxns.txt file
open(ALLRXNS, ">all_rxns.txt") or die "could not write all_rxns.tx, reason: $!\n";

#write the leading "/"
printf ALLRXNS "\/\n";

#combine the two metabolite lists
my @all_rxns = @db_rxns;

for(my $c=0; $c <= $#model_rxns; $c++) {

	push @all_rxns, $model_rxns[$c];

}  

#remove redundant metabolites
my $nr_all_rxns_ref = &RemoveRedundancies(\@all_rxns);
my @nr_all_rxns = @$nr_all_rxns_ref;

#write the non-redundante metabolite list
for(my $d=0; $d <= $#nr_all_rxns; $d++) {

	printf ALLRXNS "%s\n", $nr_all_rxns[$d];

}

#put trailing "/"
printf ALLRXNS "\/";

#create the combined Sij matrix
open(MODELSIJ, "<Sij_m.txt") or die "could not open model Sij matrix, reason: $!\n";
chomp(my @model_sij = <MODELSIJ>);
pop @model_sij;
shift @model_sij;

open(DBSIJ, "<Sij_db.txt") or die "could not open database Sij matrix, reason: $!\n";
chomp(my @db_sij = <DBSIJ>);
pop @db_sij;
shift @db_sij;

my @sij_all = @db_sij;

for(my $e = 0; $e <= $#model_sij; $e++) {

	push @sij_all, $model_sij[$e];

}

my $nr_sij_all_ref = &RemoveRedundancies(\@sij_all);
my @nr_sij_all = @$nr_sij_all_ref;

open(SIJALL, ">Sij_all.txt") or die "could not write to Sij_all.txt, reason: $!\n";

printf SIJALL "\/\n";

#create biomass precursor file
open(BIOPRE, ">biomass_precursors.txt") or die "could not create biomass precursor file, reason: $!\n";

printf BIOPRE "\/\n";

for(my $f = 0; $f <= $#nr_sij_all; $f++) {

	#print Sij line
	printf SIJALL "%s\n", $nr_sij_all[$f];
	
	#create biomass precursors files by looking for biomass reaction
	#do this by finding Sij entries where the reaction is named "biomass" then get compounds
	if($nr_sij_all[$f] =~ /\'(.+?)\'\.\'biomass\'/) {
	
		printf BIOPRE "\'%s\'\n", $1;
	
	}

}

printf SIJALL "\/";

printf BIOPRE "\/";

printf "Successfully created all OptFill input files\n\n";
```
