## Supplemental File 6 for "A Novel Optimization-Based Tool to Automate Infeasible Cycle-Free Gapfilling of Genome-Scale Metabolic Models"

Due to strict limitations on the number of supplemental files which may be included in this work, it was necessary to combine a number of supplemental text files into a single zip file which may be later retrieved and used by interested parties. Therefore, I have created this supplemental document to explain how to use the other supplemental files included in the manuscript and replicate this work and what they do.

**Files Contained in Supplemental File 3 (alphabetical order):**

**convert_XXX.py –** A set of python codes designed to take the list of reaction IDs and stoichiometries which may be found in a database or model file and create GAMS-usable files necessary to run the GAMS codes contained herein. The “XXX” portion gets replaced with the model or database name, for instance “convert_TM1.py” creates the GAMS input files for the first test model (TM1) contained in”TM1.txt” and so forth. Note that each “convert_XXX.py” file create 9 input files for GAMS. For this reason we include the code to generate these files, and not the full set of files themselves.

**FVA_combined_iJR904.gms –** GAMS code which performs FVA on the combined pseudomodel of iJR904 and iAF1260-based database used to determine which reactions were capable of holding flux. Necessary to run the “prep_for_optfill_iJR904_iAF1260.pl” code first as this code produces the necessary input for these codes.

**get_solve_times.pl –** Perl code which, by editing line 6 to any optfill output, will read the OptFill output and turn it into a handy csv (comma separated values file) which gives a much more compact view of the OptFill results such as that which can be found in Supplemental_File_5. Basically gives the results in terms of “what was the optimal values?” and “how long did they take to find?”, without all the detail of the OptFill_result_XXX_YYY.txt files.

**iAF1260_as_db.txt –** Stoichiometry file for the set of reactions from iAF1260 which remain in the final iAF1260-based database for the OptFilling of iJR904 (after having undergone the changes described in the methods). This has undergone the formatting changes of Supplemental File 1 to make the implementation easier.

**iJR904.txt –** The iJR904 model stoichiometry and reaction labels after formatting by Supplemental_File_1.

**make_m_and_db.pl –** Perl code which randomly sorts the set of new reactions which are added to the TM3/TDb3 files that are listed into those which are to be added to TM2 (to make TM3) and those to form the third test database (TDB3).

**OptFill_XXX_YYY.gms –** The GAMS code which implements the OptFilling of model XXX using database YYY. “XXX” and “YYY” are supplemented with model/database pair in the files included here. It should be noted that the “prep_for_optfill.pl” code should be run before OptFill_XXX_YYY.gms so that all necessary input files are generated for any given model/database pair. Also contains code to run FVA on each CPs solution.

**OptFill_result_XXX_YYY.gms –** These files contain the output results of a given run of “OptFill_XXX_YYY.gms” for model XXX and database YYY.

**OptFill_TFP_only_ZZZ.gms –** This is the modified OptFill code that will run only the modified TIC-Finding Problem (mTFP). This was run on only 4 models/databases, therefore “ZZZ” is substituted with “iJR904”, “iAF1260”, “TM3”, and “TDb3” only.

**prep_for­_optfill_XXX_YYY.pl –** A Perl code which runs “convert_XXX.py” and “convert_YYY.py”, then uses the outputs of the convert codes to create the input files to define the large sets. For instance, this code combines *J^M^* and *J^Db^* sets defined by the “convert_XXX.py” and “convert_YYY.py” files in the full set of reaction, *J*. This creates the files which neither convert file creates. Again, “XXX” represents the model and “YYY” represents the database in the code name which will then be substituted by the appropriate name.

**T3_all_new_rxns.txt –** The list of all the new reactions added to the TM3/TDb3 files. This is the input file for “make_m_and_db.pl” which sorts the reactions, 20% to the database, 80% to the model.

**TDb1 –** The first test database (TDb1) stoichiometry file.

**TDb2 –** The second test database (TDb2) stoichiometry file.

**TDb3 –** The third test database (TDb3) stoichiometry file.

**TM1 –** The first test model (TM1) stoichiometry file.

**TM2 –** The second test model (TM2) stoichiometry file.

**TM3 –** The third test model (TM3) stoichiometry file.

**To recreate this work using the provided code and files, using TM1/TDb1 as an example:**

1) Download/Install Perl (if on Windows, Apple iOS has Perl installed already)
2) Download/Install Python
3) Download/Install GAMS
4) Unzip Supplemental_File_3
5) Run “prep_for_optfill_TM1_TDb1.pl”
6) Run “OptFill_TM1_TDb1.gms”
7) Find your results in “OptFill_result_TM1_TDb1.txt” once the run initiated in the previous step is finished.
